# Sociality and Interaction Envelope Organize Visual Action Representations

**DOI:** 10.1101/618272

**Authors:** Leyla Tarhan, Talia Konkle

## Abstract

Humans observe a wide range of actions in their surroundings. How is the visual cortex organized to process this diverse input? Using functional neuroimaging, we measured brain responses while participants viewed short videos of everyday actions, then probed the structure in these responses using voxel-wise encoding modeling. Responses were well fit by feature spaces that capture the body parts involved in an action and the action’s targets (i.e. whether the action was directed at an object, another person, the actor, and space). Clustering analyses revealed five large-scale networks that summarized the voxel tuning: one related to social aspects of an action, and four related to the scale of the interaction envelope, ranging from fine-scale manipulations directed at objects, to large-scale whole-body movements directed at distant locations. We propose that these networks reveal the major representational joints in how actions are processed by visual regions of the brain.

**Significance Statement:** How does the brain perceive other people’s actions? Prior work has established that much of the visual cortex is active when observing others’ actions. However, this activity reflects a wide range of processes, from identifying a movement’s direction to recognizing its social content. We investigated how these diverse processes are organized within the visual cortex. We found that five networks respond during action observation: one that is involved in processing actions’ social content, and four that are involved in processing agent-object interactions and the scale of the effect that these actions have on the world (its “interaction envelope”). Based on these findings, we propose that sociality and interaction envelope size are two of the major features that organize action perception in the visual cortex.

---

We witness a multitude of actions in daily life: running, jumping, cooking, cleaning, writing, and painting, to name just a few. How does the brain make sense of this diverse input? The process begins with basic perception, forming lines and shapes into bodies in motion with identities and kinematic properties, and ultimately derives rich representations about emotional states, social interactions, and predictions about what will happen next (Kable et al., 2002; Urgesi et al. 2014; Lingnau & Downing, 2015). Much previous work on the nature of action representation in the brain has focused on the end-stages of this process; for example, attempting to localize the more abstract and conceptual aspects of action representation (e.g., Nelissen et al., 2005; Bedny et al., 2011; van Elk et al., 2013; Watson & Buxbaum, 2014; Wurm & Caramazza, 2019). However, recent research has begun to examine action-related processing from a more perceptual angle, asking how regions involved in high-level vision are organized to support action observation (Lingnau & Downing, 2015; Wurm & Caramazza, 2017). The present work follows this latter approach, leveraging an encoding analytical framework to characterize action responses in the visual system (Mitchell et al., 2008; Huth et al., 2012; Lescroart et al., 2015).

To employ this framework, we considered three core theoretical questions and their methodological implications. The first relates to the nature of the domain in question: what do we mean by “actions” and how should we sample this domain representatively (cf. Brunswick, 1955)? Many prior studies have used the tight link between actions and verbs as a guide, and have measured responses to written verbs or used verbs to guide the selection of pictures or videos of actions (Bedny et al., 2011; Daniele et al., 2013; Leshinskaya & Caramazza, 2014). However, common verbs are not always commonly-seen actions (e.g., “can”), and many commonly-seen actions are not often talked about (e.g., “chopping vegetables”). Further, using verbs (particularly without considering the type of verb) can unintentionally constrain one’s hypothesis space by implying that actions that look very different but are described by the same verb (“pushing a button” vs. “pushing a person”) share a neural representation (Hafri et al., 2017). To understand action observation at a more perceptual, rather than conceptual level, we instead sampled our action stimuli based on common human experiences, using the American Time Use Survey as a guide (see **Methods**; cf. Greene et al., 2016). These actions span a wide range of activities, such as cooking, traveling, exercising, and recreating, that a large set of Americans reported engaging in on a daily basis.

A related consideration is how best to depict these actions in order to probe underlying neural responses at a meaningful level of abstraction. Historically, researchers have relied on highly-controlled action stimuli (Jastorff et al., 2010; Isik et al., 2017), often sampling primarily from hand actions such as reaching or hammering (Watson & Buxbaum, 2014; Fabbri et al., 2016; Wurm et al., 2017). However, these well-designed and controlled stimuli are fundamentally limited in the range of actions they depict, which in turn constrains researchers’ ability to discover joints in the broader domain of action perception. To circumvent this limitation, others have taken a highly unconstrained approach, using rich, complex stimuli such as feature-length films (Huth et al., 2012; Chen et al., 2017; Huth et al., 2016). This approach comes much closer to reflecting our daily perceptual experience with actions; however, too much complexity can make the resulting data challenging to wrangle into interpretable results, especially when using thousands of features to characterize the brain’s responses to those actions. Thus, in the present work, we took an approach that sits in between these extremes: we selected a targeted and diverse subset of everyday actions, depicted using complex, heterogenous videos of a short duration (**Figure 1**).

**Figure 1.**
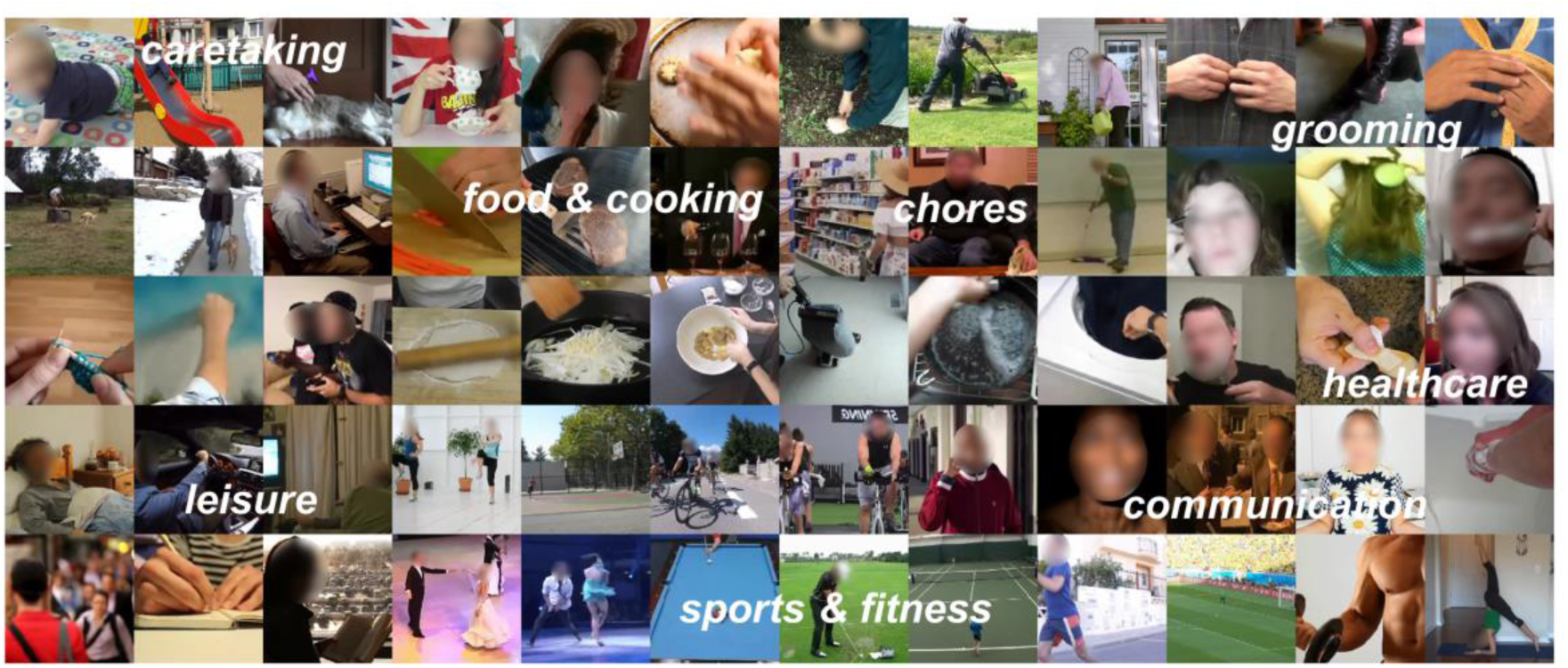
Action Video Stimuli. For each of the 60 actions in the stimulus set, a key frame is shown from one video. The key frames are arranged loosely by semantic properties for illustration purposes. These actions were selected to widely sample the space of everyday actions, based on commonly-occurring activities listed in the American Time Use Survey.

The second major consideration concerns which brain regions to target. Prior work has revealed that watching other people’s actions engages a broad network of regions with nodes in all major lobes of the brain, known as the *action observation network* (Caspers et al., 2010; Kalenine et al., 2010; Jola et al., 2012; Urgesi et al., 2014). However, the perceptual processing mechanisms that operate over objects, bodies, and motion likely also play a role in representing ecological actions. Therefore, it is critical to consider more widespread action responses across broad swathes of occipitotemporal and parietal cortices (e.g. Hafri et al., 2017; Wurm & Caramazza, 2017). Given this, we used a novel method to identify voxels that reliably differentiate among different actions (Tarhan & Konkle, under review), rather than constraining our analyses to specific regions of interest emphasized in the literature.

The final major consideration concerns the nature of the tuning in these regions. That is, what is it *about* an action that makes a given voxel respond more to that action than to others? The possibilities are potentially infinite, as actions differ from one another along many dimensions; luckily, the literature proposes several critical features and properties that may underlie action representations. For example, actions can be analyzed both in terms of the *means* by which they are performed (e.g., their kinematics and body postures) and the *ends* that they are targeted at (e.g., the actor’s intentions; Lingnau et al., 2015; Kemmerer et al., 2008). Means and ends features may be represented by distinct regions, as evidence suggests that visual responses are organized by body parts in the lateral temporal cortex (Beauchamp et al., 2002; Bracci et al., 2015), while information about an actor’s goal is processed by the inferior parietal sulcus (Hamilton & Grafton, 2006). As another example, linguistics research finds that verbs can be divided into classes based on whether they are directed at another entity (“dyadic”), and whether they entail a change of state in the world (Kemmerer et al., 2008; Levin, 1985; Levin, 1993). Relatedly, recent empirical work suggests that action processing in the lateral temporal cortex is organized by two major dimensions: ociality—whether or not an action is directed at a person -- and transitivity--whether or not it is directed at an object (Wurm et al., 2017; Lingnau & Downing, 2015).

In the present study, we operationalized some of these proposed properties, specifically the body parts that are involved in performing an action and what the action is directed at (its target; e.g., an object, another person, et cetera). Within the broad context of action processing, we consider these to be “high-level perceptual” features (as opposed to more conceptual or non-visual features). For example, we can recognize that “knitting” engages the hands and is directed at an object without any knowledge about the actor’s identity or what knitting is for. Thus, these features relate more directly to the visual processing of actions, in contrast to the more abstract properties of actions and verbs that have been the focus of much of the literature.

With these considerations in mind, the goal of this study was to understand and characterize what properties of actions best predict corresponding neural responses, allowing for the possibility that different regions of cortex are sensitive to different properties. To preview, we found that the effectors used to perform an action and what an action is targeted at successfully predicted responses to new action videos in much of the occipito-temporal and parietal cortices. Further, an analysis of the voxel-wise tunings to these features revealed a large-scale organization of five networks that span the ventral and dorsal visual streams, whose tuning is related to an action’s spatial scale of interaction and relevance to agents. We argue that these networks reflect meaningful divisions in how actions – and, more broadly, visual inputs across domains – are processed by the brain.

## RESULTS

To investigate neural responses to a wide range of human movements, 60 actions were selected based on what a large sample of Americans reported performing on a daily basis (U.S. Bureau of Labor Statistics, 2014). These actions span several broad categories such as personal care, eating and drinking, socializing, and athletics (**Figure 1**). Two short (2.5 s) videos were selected to depict each action then divided into two sets. Each video depicts a sequence of movements (e.g., stirring a pot) rather than an isolated movement (e.g., grasping a spoon, taking one step), and thus may alternatively be thought of as depicting “activities” rather than “actions.” However, “activity” can also connote something done over a much longer timescale and with greater variation in perceptual properties, such as “cooking dinner.” Thus, for clarity we refer to the stimuli as *action* videos. Participants (*N*=13) watched these action videos while undergoing functional magnetic resonance imaging (fMRI) in a condition-rich design that enabled us to extract whole-brain neural responses to each video (see **SI Appendix**).

### Voxel Selection Reveals Reliable Regions

Before analyzing the structure in these neural responses, we first asked which regions of the brain reliably differentiate the actions at all—that is, which regions systematically respond more to some actions than others. To do so, we utilized a reliability-based voxel selection method (Tarhan & Konkle, under review). This method computes split-half voxel reliability between responses to the videos in video sets 1 and 2, and then selects a cutoff that maximizes the average multi-voxel pattern reliability to each video (**Figure 2**, see **Methods**). This selection method is well-suited to our voxel-wise modeling approach for two reasons. First, it retains voxels that show systematic differences in activation across videos -- which we will ultimately try to model and predict -- removing any voxels that are either unreliable or respond equally to all actions. Second, for any action (e.g., riding a bike), the two example videos in each set differed in many respects (including the direction of movement, background, identity of the actor, et cetera). Thus, any voxels that respond similarly across these two video sets have some tolerance to very low-level features.

**Figure 2.**
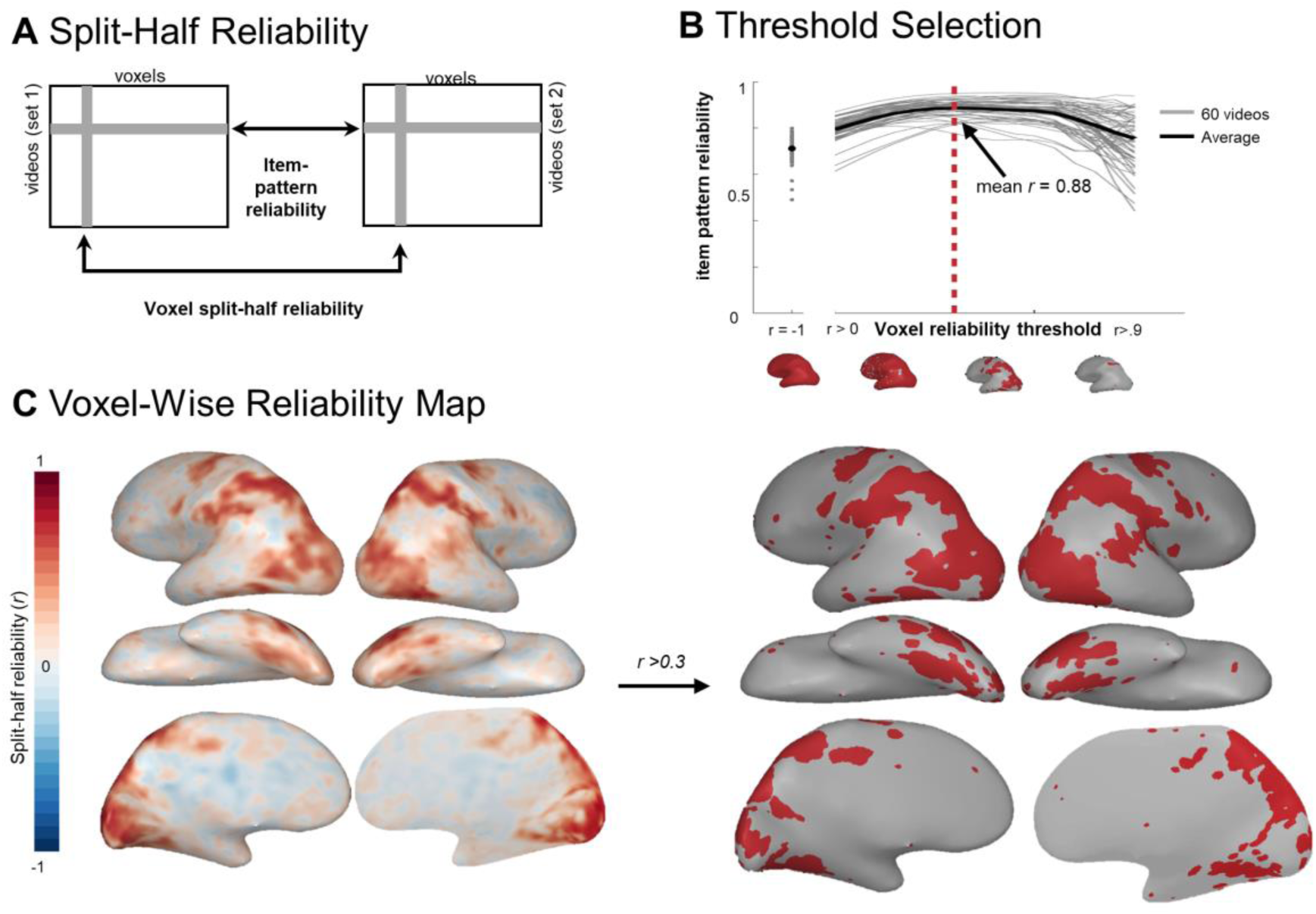
Reliability-Based Voxel Selection. A) Schematic illustrating how voxel split-half reliability and item-pattern reliability were calculated from whole-brain response data. B) Plot of average item-pattern reliability (split-half reliability of each item’s multi-voxel pattern; y-axis) among voxels that survive a range of reliability cut-offs (x-axis). Brains along the x-axis display the voxels that survive the reliability cut-offs at *r* = −1, 0, 0.35, and 0.7. C) Whole-brain map of split-half voxel reliability. D) Reliable voxels (*r*>0.30) selected based on the point where the curve plotted in (B) begins to plateau (see Tarhan & Konkle, 2019 for details). These results are based on group data. All analyses were conducted over reliable voxels, which were selected using the procedure outlined here.

The split-half reliability for each voxel is plotted in **Figure 2c** (see **Figure S2** for single-subject data). To select a cutoff for “reliable” voxels, we swept through a range of possible thresholds to find one that maximized both coverage and reliability (see **Methods**). Through this procedure, a voxel-reliability cut-off of *r*≥0.30 was selected, yielding an average item-pattern reliability of *r*=0.88 in the group data (**Figure 2b**). This method revealed reliable activations along an extensive stretch of the ventral and parietal cortices, with coverage in lateral occipito-temporal cortex (OTC), ventral OTC, and the intra-parietal sulcus (IPS) (**Figure 2c**). Reliability was relatively low in early visual areas, as expected from the cross-set reliability computation. All subsequent analyses were performed on this subset of reliable voxels.

### Operationalizing Feature Spaces with Behavioral Ratings

Among the possible features that differentiate actions, we hypothesized that the *body parts* involved in an action and *what an action is directed at* are important features that underlie at least some of the neural responses to actions. To measure the *body parts* feature space, human raters completed an online experiment in which they selected the body parts that were engaged by each action from among 20 possible effectors, such as legs, hands, eyes, torso, and individual fingers. The final feature values for each action were averaged over participants and are depicted for an example action in **Figure 3b**.

**Figure 3.**
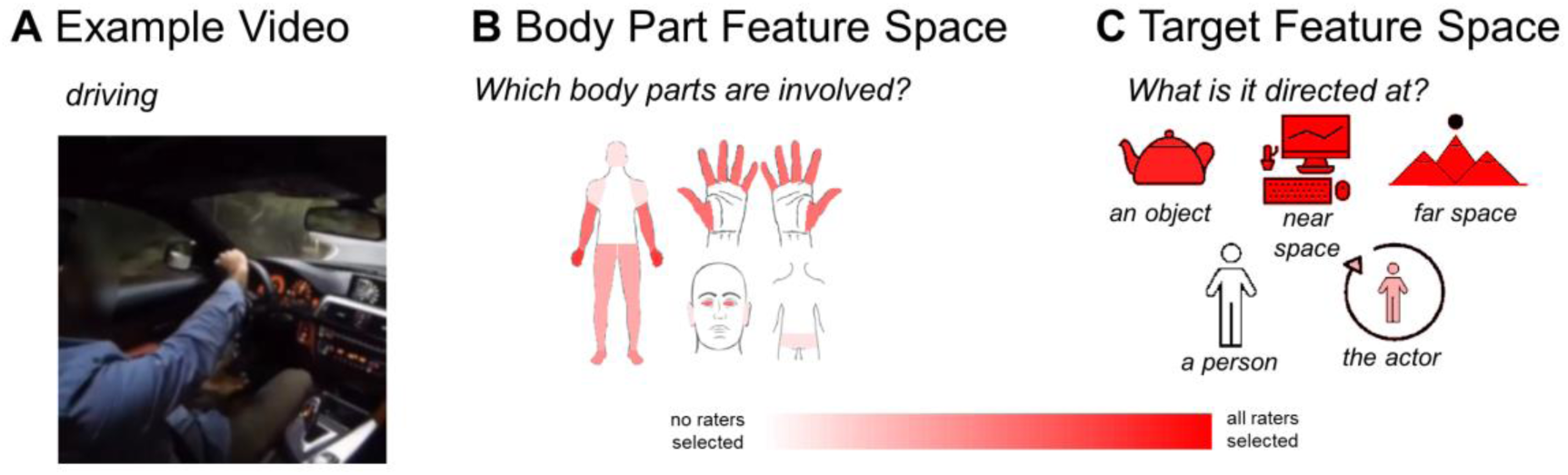
Feature Spaces. For each action video, the body parts and action targets involved in the action were estimated through online behavioral ratings experiments. (A) Keyframe from an example video depicting the action “driving.” (B) Subjects indicated which parts of the body were involved in the action video using a clickable body map. Color saturation indicates the number of participants who responded that each body part was involved in this example video. (C) Subjects rated the actions’ targets by answering the following yes-or-no questions, aimed at one of the five possible targets: object: “Is this action directed at an object or set of objects?”; near space: “Are the surfaces and space within this actor’s reach important for the action being performed?”; far space: “Is a location beyond the actor’s reach important for the action being performed?”; another person: “Is this action directed at another person (not the actor)?”; the actor: Is this action directed at the actor themselves?”. Color saturation indicates the number of participants who responded that each target was involved in this example driving video.

Given the natural covariance between different effectors, we used a principle components analysis to reduce this feature space into 7 principle components (PCs), which together account for 95% of the variance in the ratings. Inspecting how the body parts load on each PC reveals how different kinds of actions engage different groups of effectors in this stimulus set (see **Figure S1a**). For example, body part PC1 distinguishes between actions that engage the legs (e.g., running) and those that engage the hands (e.g., painting). Given that we selected actions that are commonly seen, not actions that maximally span the possible effectors, these principle components likely highlight the major body part synergies and distinctions in the space of everyday actions (e.g., perambulatory actions that center around the legs and feet versus manual actions that tend to require finer control).

Our second hypothesized feature space captures information about *action targets*, i.e. what an action is directed at. To measure these features, raters answered questions about whether the action in each video was directed at an object, another person, the actor, the reachable space, and a distant location. Actions could have multiple targets. These ratings were averaged across participants and are shown for an example video in **Figure 3c**. Following a principle components analysis, all five PCs were needed to account for 95% of the variance in the ratings (**Figure S1b**). These PCs reveal the covariance across action targets within this sample: for example, the first PC distinguishes between actions that are directed at an object (e.g., shooting a basketball) and those that are directed at another person or the actor (e.g., shaking hands or running).

Together, the seven body-part principle component dimensions and five action-target principle component dimensions were combined into a 12-dimensional feature space that was used to model brain responses to the action videos. The dimensions from the body part and action target feature spaces were not well-correlated with each other (mean *r* = 0.02, range = −0.40 – 0.45; **Figure S1c**).

### Voxel-wise Encoding Models Predict Action Responses

The primary question of this study is whether these hypothesized feature spaces can characterize neural responses to actions well, and if so where. To answer this question, we employed a voxel-wise encoding-model approach (Mitchell et al., 2008; Huth et al., 2012), which measures how well each voxel’s tuning along these feature dimensions can predict its response to a new video. Put another way, this method asks: what weighted combination of the individual features in a feature space best predicts a voxel’s responses across action videos? For example, a voxel that responds strongly to videos of sautéing and hammering but not to jumping will be best fit by high weights on hand, arm, and object-directed features, and low weights on the leg and actor-directed features. This model can then predict new responses; for example, the same voxel should respond more to knitting than to running. The quality of the model’s fit was assessed for each voxel based on how well it made these predictions for a held-out dataset (measured as a high correlation between the actual and predicted responses to the held-out data; see **Methods**).

We therefore used predictive performance to assess whether and where the body part and target feature dimensions characterize each voxel’s tuning properties well. **Figure 4 (inset)** illustrates how well the neural responses were predicted by fitting tuning weights to body part and action target features. Voxel responses were predicted well across much of the ventral and dorsal streams (median prediction across video sets (*r*_CV_) = 0.41 (set 1) & 0.42 (set 2), max *r*_CV_ = 0.76 (set 1) & 0.71 (set 2)). These patterns replicated in individual subjects, albeit at a lower overall level of prediction, which is likely related to differences in reliable voxel coverage across subjects (set1: median *r*_CV_ = 0.18, max *r*_CV_ = 0.64; set 2: median *r*_CV_ = 0.20, max *r*_CV_ = 0.67; **Figure S4**).

**Figure 4.**
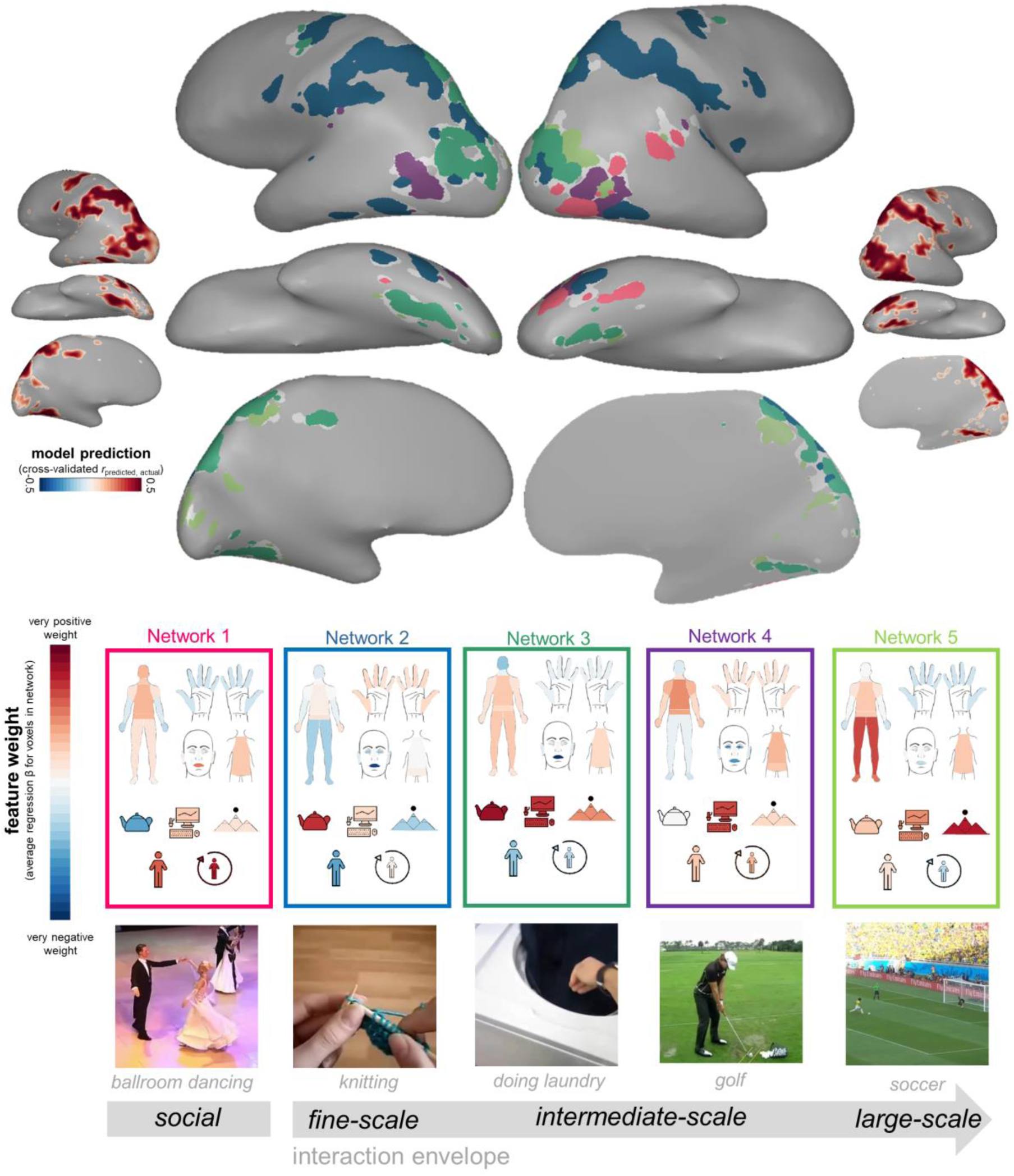
Large-Scale Feature Tuning Structure. Data-driven clustering of the feature weights for body parts and target features. Inset brains display the prediction performance (*r*_CV_, the correlation between predicted and actual responses in held-out data) for the combination of these features within reliable voxels. Larger brains display the five large-scale clusters on an example subject’s brain. Heat maps display each cluster centroid’s feature weight profile. Key frame images depict example videos that elicited high responses from each network. Results are displayed for video set 1 (see also **Figure S3**). Only voxels that were well-fit by the model in both video sets (positive *r*_cv_ that was significant after FDR correction for multiple comparisons) were analyzed. Voxels surviving this criterion with insufficient variance to be included in the clustering analysis are colored grey.

Critically, the key advantage of this encoding model framework is that we can examine not just *how well* the model predicts a voxel’s response to a new action, but also *why*. For example, some voxels might be tuned to leg and foot involvement, while others might be tuned to hand and mouth involvement. To understand how these feature tunings are mapped across the cortex, we used *k*-means clustering to group voxels by their model coefficients (see **Methods**), considering only voxels whose responses were fit reasonably well by the model (*r*_CV_>0 and significant after corrections for multiple comparisons). This analysis clusters voxels together if they have similar feature tunings (e.g., voxels assigned high weights on hand involvement and object targets but not leg involvement will be grouped together). Importantly, this method does not require that voxels are grouped into contiguous clusters, making it possible to discover both contiguous regions and networks of non-contiguous regions that have similar tuning functions. Further, this method does not presuppose any particular combinations of feature tunings ahead of time, allowing natural patterns to be revealed directly from the data (see Webster & Fine, in prep; Lashkari et al., 2010; Vul et al., 2012 for related analysis approaches). Through this method, we found evidence for five networks that are tuned to different characteristics of actions. We chose this five-network solution by considering silhouette distance and cluster center similarity (see **Methods, Figure S3a**). Each network is shown with its corresponding tuning function in **Figure 4**.

Network 1 (pink) is primarily right-lateralized, covering regions along the fusiform gyrus and extending between the occipital face area (OFA) and superior temporal sulcus (STS). This network is tuned to face features and not hands and is directed at people (others and the actor) but not objects. It is near STS regions typically engaged by social processing (Saxe et al., 2004; Deen et al., 2015; Isik et al., 2017) while also extending inferiorly into ventral OTC along the fusiform gyrus. This network’s tuning pattern highlights a possible large-scale neural division between social or body-centric actions that are directed at other people (shaking hands) or the actor themselves (laughing), and those directed at objects and space. In fact, this division emerges early on: Network 1 separates out from the rest when voxels are grouped into just two networks (**Figure S5**), a finding we confirmed using hierarchical clustering (**Figure S6**; see **SI Appendix**). This pattern suggests that the distinction between social or agent-focused actions and nonsocial actions is the predominant joint organizing action responses in these regions.

The remaining four networks are tuned to non-social aspects of action. The dorsal stream responses divided into two prominent networks. Network 2 (blue) contains voxels that stretch extensively along superior IPS, as well as a “satellite” node in lateral OTC. Inspecting the feature weights associated with this network indicates that these voxels respond most strongly to videos that involve finger and hand movements, and that are directed at objects in the near space (e.g., knitting). This finding resonates with a previously-established tool network (Johnson-Frey et al., 2004; Hasson et al., 2004; Grafton et al., 2007; Kalenine et al., 2010; Lingnau & Downing, 2015; Baldassano et al., 2016; Wurm et al., 2017). Network 3 (dark green) contains voxels that stretch extensively along inferior IPS, into the transverse occipital sulcus (TOS), with a satellite node in the parahippocampal cortex (PHC). Inspecting the feature weights associated with this network also reveals strong tuning to both objects and near space, but with more arm (and not finger) involvement.

Network 4 (purple) is restricted to the ventral stream, including bilateral regions in the vicinity of extrastriate body area (EBA). Regions in this network are tuned to hands, arms, the torso, and near space.

Finally, Network 5 (light green) has nodes along PHC, TOS, and the medial surface of the cortex in both the parietal and retrosplenial regions. This final network effectively runs parallel to Network 3 (dark green): both networks anatomically resemble scene-preferring regions, such as the parahippocampal place area, occipital place area, and retrosplenial cortex (Epstein & Kanwisher, 1998; Epstein, 2005). However, compared to Network 3, Network 5 is tuned more strongly to actions directed at far space, the legs, and the whole body and less strongly to actions directed at objects. These differences in tuning echo prior work showing that at least one scene-preferring region – PPA – contains sub-regions for object and space processing (Baldassano et al., 2016).

Inspecting these last four non-social networks reveals a possible overarching organization, where each network has a preference for a different *interaction envelope*, or the scale of space at which that agent-object interactions take place in the world. The four interaction envelopes move from small-scale, precise movements involving hands and objects (knitting), to less precise movements also involving hands and objects (loading a washing machine), to intermediate-scale movements involving the upper body and near spaces (golfing), to large-scale movements that engage the whole body and require more space (playing soccer).

To examine the robustness of these findings in video set 1, we also conducted the same analysis on voxel tuning weights fit using video set 2. We then measured the correspondence between the two sets’ network solutions using d-prime, a signal-detection measure that captures whether voxels were assigned to the same networks for both sets. This five-network solution was quite consistent across the two video sets (cross-sets d-prime: 1.6; **Figure S3b & c**), pointing to the robustness of this network structure. In general, we also found that the single-subject data echoed these patterns (**Figure S4**). For example, we found a cluster in the right superior temporal sulcus that is tuned to people in 9/13 subjects. In addition, we found that 10/13 subjects had a pair of clusters that spanned the intra-parietal sulcus and lateral OTC and were tuned to fine- and intermediate scales of space (analogous to Networks 2 and 3 in the group data). Further, 9/13 subjects had 3-4 networks whose tuning ranged from fine, to intermediate, to large scales of space, just as we observed in Networks 2-5 in the group data. However, not all subjects reflected the group data perfectly. In general, subjects with more extensive reliable coverage were a better reflection of the group, while those with fewer reliable voxels showed greater variability.

Taken together, these voxel-wise modeling and clustering analyses provide evidence for five networks that distinguish predominantly between social and non-social actions, then further divide the non-social actions into four large classes that vary in the scale of space at which they affect the world. These data-driven analyses reveal that the relevance to agents and scale of an action in the world may be critical organizing factors for the perception and representation of actions in the brain.

### Convergence between feature-based and response-based networks

To arrive at the conclusion that there are 5 major subnetworks underlying visual action perception, we relied on our hypothesized feature spaces, which characterize actions’ body part involvement and targets. However, it is also possible that there is more systematic structure in the responses to these actions that is not captured by these feature spaces. In this case, the five networks we find may be only a partial reflection of the true sub-networks patterning this cortex.

To examine this possibility, we performed another variant of the *k*-means clustering analysis, this time grouping voxels based on their profile of responses to all 60 videos in a stimulus set, rather than the profile of their feature-tuning weights. Thus, the resulting clusters are driven entirely by the brain’s responses to the videos themselves; the body part and target features did not enter into the analysis. The results of this analysis are displayed in **Figure 5**. We again found evidence for five networks (cross-sets d-prime: 1.7) that recover a relatively similar structure to that observed based on the feature tuning (d-prime between feature-based and feature-free solutions: 1.6 for both video sets). The convergence between the results of the feature-free and feature-based analyses provides empirical evidence that the large-scale structure described above is not merely an artifact of the features we chose (see also **Figure S7**).

**Figure 5:**
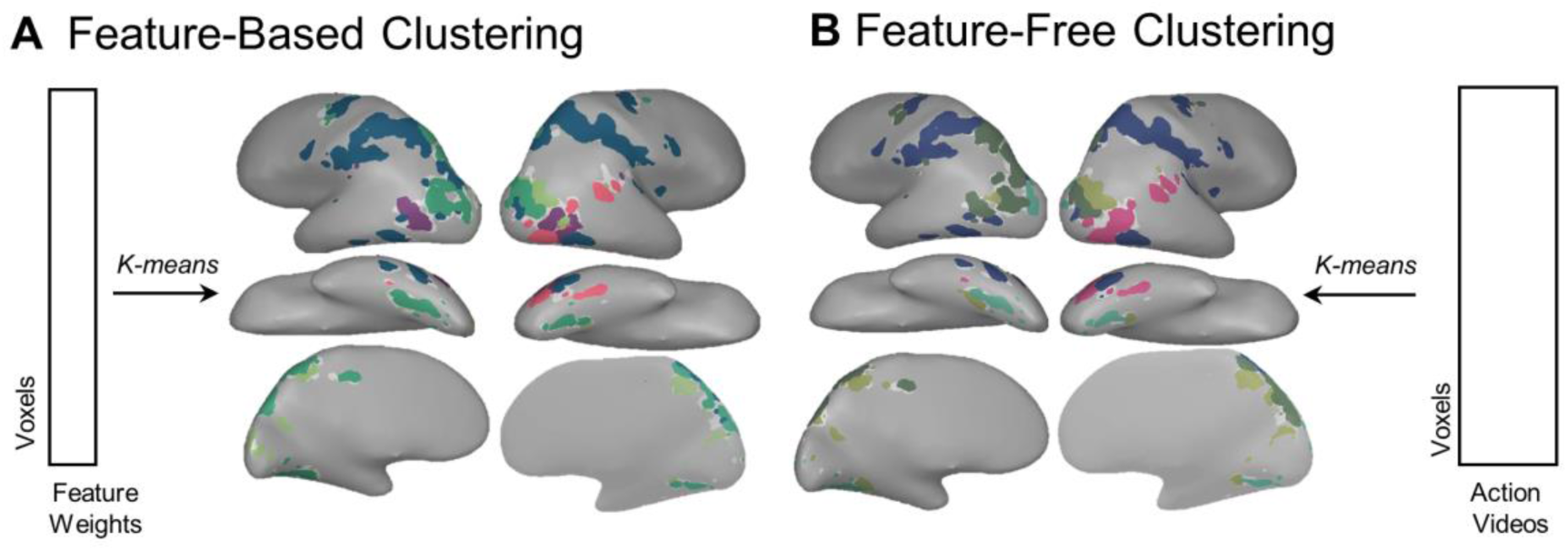
Generality of the Large-Scale Structure. We compared the results of clustering voxels based on the feature weights fit by the encoding models (A) and by the raw activation patterns across all 60 videos (B). The 5-network solutions over data from video set 1 are shown for both clustering analyses. Parts of cortex with similar colors had either similar feature weights (A) or similar overall response profiles (B). Each cluster’s color was assigned algorithmically such that clusters with similar response profiles were similar in hue and were determined separately for (A) and (B).

### Auxiliary Questions

We next ask a series of more targeted questions to clarify the implications of these results and link them directly to related work.

#### What is the role of low-level features?

In addition to higher-level perceptual features related to an action’s target and kinematics, the brain must also process low- and mid-level features like motion and form information. For example, videos vary in whether the actor is on the left or the right of the frame, and generally each video has a different spatial distribution of visual information across the frames. How do these lower-level aspects of the videos’ visual structure fit into the neural feature space, and which regions are more strongly tuned to them?

To investigate this question, we used an image statistics model (Gist; Oliva & Torralba, 2001), to capture the spatial frequency and orientation content present in a coarse grid over each video’s frame, averaged over time (see **Methods**). We then compared the gist model’s predictive performance with that of the body-part-action-target model using a preference-mapping method: each voxel was colored according to the model that predicted its responses best. Note these two feature spaces were not well-correlated—the representational dissimilarity matrix correlation between gist and high-level features was r = 0.01 for video set 1 r= 0.05 for video set 2. Because we anticipated that low-level gist features would best predict responses in more retinotopic early visual regions, here we examined a larger set of reliable voxels, based on a variant of the voxel selection procedure that selects voxels that respond reliably *within* rather than *across* video sets and provides better coverage of early visual regions (see **Methods**).

**Figure 6** shows how these models perform across these regions. As expected, the gist features predict noticeably better in early visual cortex, but the body part and target features predominate in the temporal and parietal cortex. It is also worth noting that, even though the body part and target features largely outperform the gist features outside of early visual cortex, gist features also predicted neural responses moderately well throughout both the dorsal and ventral streams (**Figure 6a**). These results suggest that the action representations across this cortex retain information about the spatial layout of the action scene, even as higher-level features begin to emerge, echoing similar findings related to object processing (Long et al., 2018).

**Figure 6:**
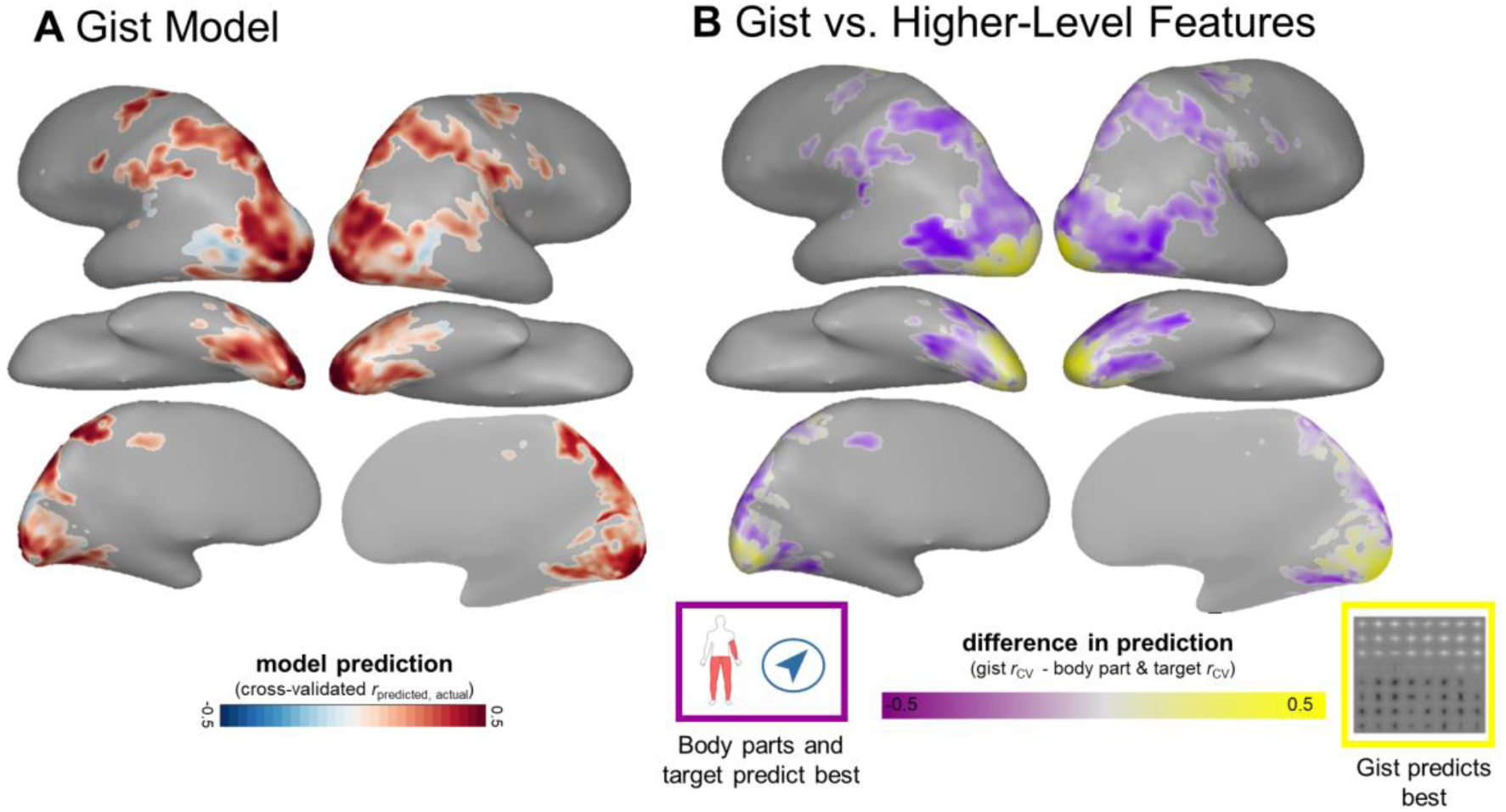
Incorporating Low-Level Visual Features. (A) Voxel-wise encoding model prediction results for the gist model in reliable voxels. Voxel color reflects the cross-validated correlation between predicted and actual response patterns to held-out items. (B) Two-way preference map comparing prediction performance for the gist features with performance for the body parts and action target features. Voxels are colored according to the model with the best cross-validated prediction performance: yellow for gist, and purple for the body parts and action target features. Color saturation reflects the strength of the voxel’s preference (*r*_CV_ for the gist model – *r*_CV_ for the body parts and action target model).

The effects of low-level motion features were not investigated in-depth in this study. It is likely that motion plays an important role in action processing (Johansson, 1973; Isik et al., 2017). However, prior work has found that motion models such as motion energy (Nishimoto et al., 2011) are only effective at capturing brain activity when subjects are required to maintain central fixation. When they view videos in a naturalistic fashion, moving their eyes around the frame as in the current study, this model breaks down unless neuroimaging is coupled with eye-tracking (Nishimoto et al. 2011, Nishimoto & Gallant 2011). Further, our use of fMRI makes it challenging to collect measurements at the fine temporal scale at which motion features vary in our short video clips. Therefore, a combination of eye-tracking and fast fMRI sampling would be necessary to understand the how motion features contribute to the large-scale organization of action processing in the brain.

A related possibility concerning motion is that the tuning of the four interaction envelope networks may reflect the span of movement in the videos; i.e., how much of a video’s frame contains motion? To investigate this question *post-hoc* we measured the span of movement in each video using two methods. First, we collected ratings of how much the average actor moves when performing each action. Second, we used an optical flow algorithm (Horn & Schunck, 1981) to measure the proportion of each video’s frame that contained any movement. We then calculated each network’s sensitivity to variations in the motion span (see **SI Appendix**). We found that motion span sensitivity tracks with the size of the interaction envelope for small and intermediate interaction envelopes, but not the largest (**Figure S8**). This pattern replicated across both motion span measures. Broadly, this line of inquiry raises further questions about what lower-level visual features predict the size of an action’s interaction envelope, which will be fruitful ground for future research.

#### Do these neural responses simply reflect what’s visible in each video?

Up to this point, we have implicitly assumed that neural responses fit by the body part and target features reflect their *involvement*. However, it is also possible that these responses reflect the features at a more primitive level: whether they are simply *visible*. These two interpretations are necessarily related (e.g., it’s difficult to naturalistically depict “knitting” without also showing the hands, though whether or not the legs are depicted in the video is less constrained by this action).

To quantify the relationship between the involvement and visibility of these features in this stimulus set, we separately measured which body parts and targets were visible in each video (see **Methods**). We found that ratings of the visible and involved features were only moderately correlated (for body parts: *r* = 0.38; for action targets: *r* = 0.30; combined feature space *r*=0.44). **Figure 7a** shows the features that differed the most between ratings of their visibility and involvement, which include the head, torso, face, and far space. Often, the body parts and targets that were rated as involved were a subset of the visible features. For example, in our video depicting the action of reading a book, the actor’s whole upper body, a reachable desk surface, and a distant scene through a window are visible; however, this action only involves the book and the actor’s eyes, hands, and arms. In other cases, body parts were rated as involved even if they were off-screen. For example, raters responded that our video depicting vacuuming, which only shows the actor’s legs and the vacuum, also involved the arms, hands, and eyes.

**Figure 7:**
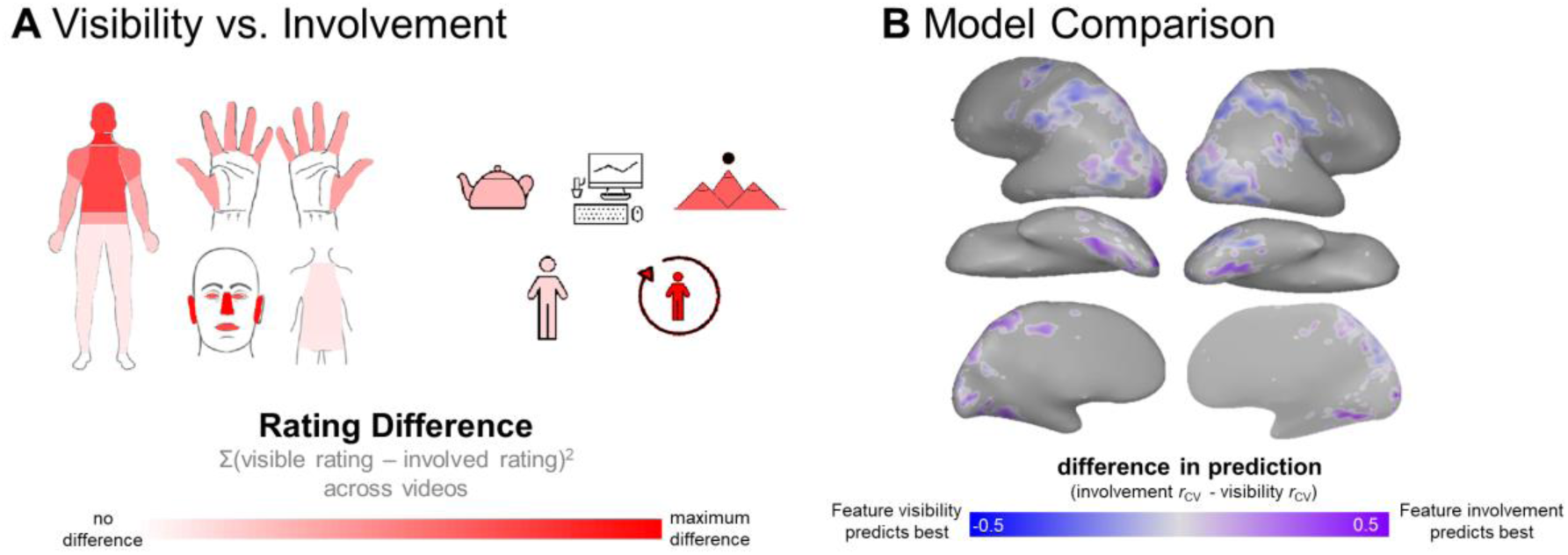
The Role of Feature Visibility. (A) Summary of feature-based differences between ratings of the features’ visibility and involvement in the actions. Body parts and action targets that dissociate most strongly between ratings of visibility and involvement across our stimulus set are colored with the strongest saturation. (B) Preference map comparing prediction performance for the models based on feature involvement and visibility. Voxels are colored according to the model with the best cross-validated prediction performance: purple for the features’ involvement, and blue for their visibility. Color saturation reflects the strength of the preference (*r*_CV_ for the involvement model – *r*_CV_ for the visibility model).

To determine whether and where the visibility or involvement of these features best predicted brain responses, we separately fit each model and then calculated which model predicted responses best in each voxel (see **Methods**). **Figure 7b** shows the results of this analysis. The model based on the features *involved* in the actions outperforms the model based on the features that are *visible* in the videos in the posterior lateral occipital cortex, lateral temporal cortex in the vicinity of EBA, and the fusiform gyrus. In contrast, and somewhat surprisingly, the model based on the *visible* features predicts better in much of the IPS. Note that this analysis does not reveal how much variance is unique to each model; in fact, we found that a standard variance partitioning analysis (e.g. Lescroart et al., 2015) produced unstable results and was therefore ill-suited to our data. Instead, this analysis indicates that the visible bodies, people, objects, and backgrounds play an important role in how these actions are processed in some regions. However, the functional involvement of these effectors and targets is a better description of the neural responses throughout the ventral stream.

## GENERAL DISCUSSION

Recognizing the actions of others is an essential capacity of the human visual system. While extensive research has examined this capacity within the motor and language systems, here we focused on the high-level perceptual properties of everyday actions. We found that much of the occipito-temporal and parietal cortices responded reliably during action observation, and these responses were well-predicted by models based on the body parts engaged by the actions and their targets in the world.

Examining the structure in voxels’ tunings to these features revealed evidence for five action-processing sub-networks. The first of these networks is tuned to agent-directed actions, including actions directed at other people and the actor themselves. The remaining four networks vary in the scope of their *interaction envelope* – the spatial extent of interactions between agents, objects, and the environment. We propose that these five sub-networks reflect deeper joints within action processing in the visual system. In the following sections, we relate these findings to prior work, and then discuss how this organization for action representations might help to better understand the broader principles organizing the visual system.

### Links to Prior Work

The five sub-networks found in our data converge with several known networks of the ventral and dorsal stream, including those that display preferences for bodies, faces, objects, and scenes (Kanwisher et al., 1997; Epstein & Kanwisher, 1998; Grill-Spector et al., 1999; Downing et al., 2001). For example, the agent-focused sub-network (Network 1) encompasses regions which are traditionally thought to be specialized for body parts, whole bodies, and faces (Downing et al., 2001; Saxe et al., 2004; Bracci et al., 2011; Grill-Spector & Weiner, 2014; Isik et al., 2017), and it is right-lateralized, consistent with prior results (e.g. Grossman & Blake, 2001; Grossman & Blake, 2002). Our findings also converge with previous findings that responses in occipito-temporal cortex are organized by animacy and object size (Konkle & Oliva, 2012; Konkle & Caramazza, 2013; Long et al., 2018). In general, big objects are processed in regions tuned to larger-scale actions, small objects are processing in regions tuned to smaller-scale actions, and animals are processed in regions tuned to agents. Additionally, the network related to large-scale interactions such as locomotion (Network 5) encompasses well-known scene-selective regions near occipital place area, retrosplenial cortex, and along the parahippocampal place area. Importantly, both face and scene networks are typically identified by measuring responses to views of isolated scenes or faces, and then contrasting these against each other or to views of objects. However, in our stimuli, bodies, objects, and scene elements appear in *all* videos. Therefore, our results likely converge with prior findings not simply because of the mere presence of these visual categories but because scenes and bodies are more relevant to some actions than others.

In some cases, our results diverge from prior work. For example, the sub-network related to large-scale actions and far spaces (Network 5), is primarily right-lateralized in both video sets. However, most research on scene networks shows relatively clear bilateral networks (Epstein & Kanwisher, 1998; Epstein et al., 2005; Epstein, 2008). Relatedly, the sub-networks tuned to object targets within a more focal interaction envelope (Networks 2 and 3) are bilateral in this data set, which is consistent with some tool-network studies (e.g. Kellenbach et al., 2003; Bracci & Peelen, 2013; Chen et al., 2017), but not others (Johnson-Frey et al., 2004; Johnson-Frey, 2004; Frey, 2008). One interesting possibility is that studying responses to actions and actors together with tools and scenes will help shed light on when and why we see hemispheric differences along the ventral and dorsal streams.

Finally, perhaps the most surprising result was that ventral stream responses were predicted best by whether body parts and action targets were *involved* in the action, while dorsal stream responses were predicted best by their *visibility* in the video. While it is clear that viewing actions drives both the ventral and dorsal visual streams, it is also clear that more work is needed to understand how the streams relate to one another during action observation. One potential avenue for clarifying this relationship is to characterize the relative contributions of what is *visible* and what participants *fixate* on. It is possible that participants fixate on the features that are more functionally-relevant to an action. If this is the case, the visual system will give these fixated features priority, and therefore responses in the ventral stream may really reflect what participants look at, which also happen to be the features that are functionally relevant to an action. It is possible that the same is not true in the dorsal stream because it represents the full action scene, regardless of what participants look at the most. Future eye-tracking studies are needed to decisively test these hypotheses.

In addition to converging with prior work on other categories of visual input, our methods build on the use of encoding models to predict responses to rich videos (Huth et al., 2012; Huth et al., 2016). However, our approaches differ in the granularity at which we predict voxel responses and subsequently infer voxel tuning properties. In Huth et al., (2012), a voxel’s response could be fit by putting weights on over 1,000 predictors, including verbs like “cooking”, “talking”, and “crawl”, as well as nouns like “tortoise” and “vascular plant.” It is possible to map some of these specific features onto our more-general ones -- for example, a voxel in their data set with high weights on “knitting” and “writing” might be fit in our data set by high weights on the hand, fingers, and object-target features. However, one potential advantage of characterizing the brain’s responses at the level of body parts and targets is that the features are more generative: it is simple to map any new action into this low-dimensional feature space, and this level of representation may therefore be more appropriate for characterizing the response tuning of mid-to-high-level visual cortex.

### Sub-Networks for Action Perception

One strength of the current approach is that we considered neural responses to a widely- and systematically-sampled set of actions. In doing so, we found that the most predominant division present in neural responses was between actions focused on the agents and those focused on objects and space (**Figure 4; Figure S5, Figure S6**). That is, viewing actions directed at the self or others seems to recruit some regions of the visual system to a different degree than viewing actions directed at objects and space. In some respects, this result is broadly consistent with Wurm & Caramazza (2017)’s proposal that sociality is an organizing property of action representation. However, we did not find converging evidence for their anatomical proposal that tuning for actions’ sociality and transitivity is reflected in a ventral-dorsal organization across the lateral OTC (see **SI Appendix, Figure S9**). Rather, we found that the dorsal stream and parahippocampal gyrus are more sensitive to object-directed actions, while most of lateral OTC and the fusiform gyrus are sensitive to whether an action is directed at the actor or another person. More generally, our data-driven finding of a neural joint between agent-focused and non-agent-focused actions dovetails nicely with the broader argument that humans recruit fundamentally different cognitive architectures when processing agents (e.g., inferring beliefs and goals or detecting interactions between agents) and interactions that are less focused on the agents (e.g., reaching, grasping, and navigation; Johnson et al., 1998; Sweetenham et al., 1998; Papeo et al., 2017; Isik et al., 2017).

We also found that regions tuned to non-agent-focused actions branched into four networks. We propose that these highlight different *interaction envelopes*, or the scales at which actions affect the world. These range from fine-motor, hand-focused actions like knitting, to coarser movements like doing laundry, to intermediate actions involving the upper body and near spaces like golfing, to large-scale actions that move the whole body within a larger space, like playing soccer. To our knowledge, the term “interaction envelope” was first introduced in the visual cognitive neuroscience literature to highlight the difference between objects typically used with either one hand or two (Bainbridge & Oliva, 2015). Here we have adopted this term and expanded its scope. While the scale of the interaction envelope is a fairly intuitive continuum along which to organize actions, it is also a relatively novel theoretical proposal, contrasting with the more linguistic properties emphasized in the literature, such as transitivity and communicativeness (Grafton & Hamilton, 2007; Van Elk et al., 2014; Lingnau & Downing, 2015). Given this novelty, it’s important to consider why the size of the interaction envelope might be a useful principle for organizing brain responses to observed actions.

One interesting possibility is that the interaction envelope might be useful for generating meaningful higher-level predictions about an actor. For example, a person performing a small-scale, object-directed action, like writing, likely has a focal attentional spotlight, a still body, and may be mentally but not physically taxed. At the other extreme, a person performing a large-scale action, like running, likely has a wide attentional spotlight, a dynamic body, and may be physically but not mentally taxed. Further, grouping visual action representations in this fashion resonates with important distinctions between high-level semantic categories. For example, most tool-use actions are also hand- and object-based, while fitness actions tend to engage the whole body within a larger spatial envelope. Following this logic, it may be productive to think of action processing as a form of complex scene processing that integrates object, scene, and body processing. That is, perhaps actions are a chassis that connects the visual processing of objects, bodies, and space, rather than reflecting a separate domain of visual input.

## METHODS

### Experimental Model and Subject Details

#### Human Subjects

Thirteen healthy, right-handed subjects (5 males, age: 21 – 39 years) with normal or corrected-to-normal vision were recruited through the Department of Psychology at Harvard University and participated in a 2-hour neuroimaging experiment. In addition, 802 participants completed behavioral rating experiments conducted online. All subjects gave informed consent according to procedures approved by the Harvard University Internal Review Board.

### Method Details

#### Stimulus Set

120 2.5-second videos of 60 everyday actions were collected from YouTube, Vine, the Human Movement Database (Kuehne et al., 2011), and the University of Central Florida’s Action Recognition Data Set (Soomro, Zamir, & Shah, 2012). These were divided into 2 sets of 60 videos each, so that each set contained one exemplar depicting each of the 60 actions. All videos were cropped to a 512-by-512 px frame centered on the action, with no visible logos or borders, and stripped of sound. The 60 actions were selected based on the American Time Use Survey corpus (U.S. Bureau of Labor Statistics, 2014), which records the activities that Americans perform on a regular basis, across a range of general categories (e.g., household chores, fitness, and work tasks). These categories were used to ensure that we sampled a wide range of everyday visual experience but were not used in the analyses. Key frames from both sets of videos are available for download from the Open Science Framework.

#### fMRI Data Collection

Participants viewed 120 videos of everyday actions in a condition-rich design while undergoing functional neuroimaging (**SI Appendix**). To ensure that they remained alert throughout the experiment, participants pressed a button whenever a red frame appeared around a video.

#### Action Feature Ratings

To collect action feature ratings, four behavioral experiments were conducted on Amazon Mechanical Turk. In all experiments, 9-12 raters viewed each video, named the action depicted, and answered experiment-specific follow-up questions. In Experiment 1 (*N* = 182), raters indicated the body parts that were engaged by the action in the video by selecting them from a clickable map of the human body: when a body part was selected, it was highlighted in red (**Figure 3b**). There were 20 possible body parts: eyes, nose, mouth, ears, head, neck, shoulders, torso, back, arms, hands, individual fingers, waist, butt, legs, and feet. The lateralization of body parts was not analyzed: e.g., a rating was recorded for “hand” if raters selected the right hand, left hand, or both hands. In Experiment 2 (*N* = 240), raters indicated the body parts that were visible at any point in the video using the same method. In Experiment 3 (*N* = 180), raters indicated what each action was directed at by answering five yes-or-no questions: “is this action directed at an object or set of objects?”; “is this action directed at another person (not the actor)?”; is this action directed at the actor themselves?”; “are the surfaces and space within the actor’s reach important for the action being performed?”; “is a location beyond the actor’s reach important for the action being performed?” (**Figure 3c**). In Experiment 4 (*N* = 200), raters responded to similar questions about the targets that were visible in the videos (e.g., “is an object or set of objects visible in the video?”).

#### Gist Features

For comparison with the body parts and action target features, the “gist” model (Oliva & Torralba, 2001) was included in the analysis as a measure of low-level visual variability between videos. First, each video frame was divided into an 8 × 8 grid. At each grid location we quantified the power at 4 different spatial frequencies and scales (12 orientations at the finest scale, 8 at the intermediate scale, and 6 at the coarsest scale), yielding a 1,920-dimensional feature vector for each frame. These gist features were extracted for each video frame and then submitted to a principle components analysis (PCA) to obtain features that describe the global shape of the scene. Following Oliva & Torralba (2001), the first 20 principle components were then averaged across frames in a given video, resulting in a 120-by-20 gist feature matrix. Together, these 20 principle components accounted for nearly 100% of the variance in the videos.

### Quantification and Statistical Analysis

#### fMRI Reliability and Voxel Selection

Split-half reliability was calculated for each voxel by correlating the betas extracted from odd and even runs of the main task (**Figure 2a**). This was done in two ways. Reliability was calculated *across* sets by correlating odd and even betas from glms calculated over the two video sets. Across-sets reliable voxels did not extensively cover early visual cortex, as this scheme requires responses to generalize over two different exemplars of the same action. Reliability was also calculated *within* sets by correlating odd and even betas separately for each set, then averaging the resulting *r*-maps. Within-set reliable voxels had better coverage of early visual cortex and were only used to compare the gist model to the body-part-action-target model (**Figure 6b**). For both types of reliability, we selected a reliability-based cutoff using a procedure from Tarhan & Konkle (2019), which strikes a balance between selecting a relatively small set of voxels with the highest reliability and selecting a larger number of voxels but with lower reliability. First, we plotted the reliability of each video’s multi-voxel response pattern (*item-pattern reliability*) across a range of candidate cutoffs. Then, we selected the cutoff based on where the multi-voxel pattern reliability begins to plateau for all videos. This method takes advantage of the fact that after a certain point, using a stricter cutoff restricts coverage without significantly increasing the data’s reliability. Using this approach, we determined that any voxel with an average reliability of 0.3 or higher was a reasonable cutoff for inclusion in the feature modeling analysis because it maximized reliability without sacrificing too much coverage (**Figure 2b & 2c**). This cutoff held in both group and single-subject data; however, only voxels that were reliable at the group level were analyzed.

#### Voxel-wise Encoding Models

##### Model Fitting and Validation

We used an encoding-model approach (Mitchell et al., 2008; Huth et al., 2012) to model each voxel’s response magnitude for each action video as a weighted sum of the elements in the video’s feature vector (e.g., individual body parts) using L2 (“ridge”) regularized regression. Models were fit separately for the two video sets. The regularization coefficient (λ) in each voxel was selected from 100 possible values (ranging from 0 to the maximum value that produced a non-null model for that voxel). The final λ was the value that minimized the mean-squared error of the fit in a 10-fold cross-validation procedure, using the data from the other video set. To ensure that our models were not over-fit, we estimated their ability to predict out of sample using a cross-sets cross-validation procedure. This was done by training the model in every voxel, using the data from all 60 videos in one video set. We then calculated the predicted response magnitude for the second video set (beta weights from the training model * feature vector for the held-out video set). Finally, the predicted and actual data for the held-out actions were correlated to produce a single cross-validated *r*-value (*r*_CV_) for each voxel. All models were fit using responses from the group data (see **Supplement** for single-subject results).

#### Data-Driven Neural Clustering

##### Feature-Based Clustering

We used *k*-means clustering to group voxels based on their feature weight profiles. This analysis groups voxels based on the similarity of the weights that the voxel-wise encoding model assigned to the 12 body-part and action-target features. These feature weights were measured by fitting the model on the complete dataset for each video set, and the clustering analysis was conducted separately in both Set 1 and Set 2, enabling a internal replication of the main result. In order to facilitate a comparison between the sets, we ensured these clustering analyses were conducted over the same set of voxels—thus, we only included voxels that were reliable across both video sets and were predicted well (*r*_CV_>0, with q < 0.01 after FDR-correction) by both the models. Next, we used MATLAB’s implementation of the *k*-means algorithm with the correlation distance metric to cluster voxels by feature weight profile similarity (10 replicates, 500 max iterations). The correlation metric was chosen in order to group voxels together that have similar relative weightings across the features (e.g. higher for leg involvement and lower for near space targets). For any clustering solution, the cluster centroid weight profiles reflect the average normalized profile for all voxels included in the cluster.

To determine the number of clusters to group the voxels into (*k*), we iteratively performed *k*-means on the data from video set 1, varying the possible *k* value from two to twenty (**Figure S3**). When choosing the final *k*, we considered both silhouette distance (how close each voxel is in the 12-dimensional feature-space to other voxels in its cluster, relative to other nearby clusters) and cluster center similarity (how similar the cluster centers are to one another on average). The logic for the latter measure is that if two clusters have very similar centroids, there is very little we can do to interpret their differences; thus, by considering solutions with lower cluster center similarity we are better equipped to interpret divisions within this feature space. In addition, we visualized the solutions at *k* = 2, 3, and 4 (**Figure S5**), to provide insight into the hierarchical structure of these networks.

To visualize the final clustering solution, we created cortical maps in which all voxels assigned to the same cluster were colored the same. For visualization purposes, we chose these colors algorithmically such that clusters with similar response profiles were similar in hue. To do so, we submitted the clusters’ feature weight profiles to a multi-dimensional scaling algorithm using the correlation distance metric, placing similar cluster centroids nearby in a three-dimensional space. The 3D coordinates of each point were re-scaled to fall in the range [0 1], and then used as the Red-Green-Blue color channels for the cluster color. We then plotted each cluster’s centroid to determine how to interpret the groupings. This was done by multiplying the centroid’s feature weight profile over the 12 feature PCs by the factor loading matrix from the PCA, effectively projecting the centroid back into the original feature space. The resulting weights over the 25 original features were visualized using custom icons, where each feature’s color reflected the weighting of its associated PC feature in the centroid.

As a test of robustness, we calculated the sensitivity index (d’) between the solutions for video sets 1 and 2. To do so, we created a voxels x voxels matrix for both video sets, with values equal to 1 if the voxels were assigned to the same cluster and 0 if they were assigned to different clusters. Hit rate was calculated as the percent of voxel-voxel pairs that were assigned to the same cluster in set 1 that were also assigned to the same cluster in set 2. False alarm rate was calculated as the percent of voxel-voxel pairs that were not assigned to the same cluster in set 1 but were assigned to the same cluster in set 2. Sensitivity (d’) was subsequently calculated as z(Hit)-z(FA). D’ was also calculated between the shuffled data for sets 1 and 2, producing a baseline value close to zero (mean across *k*-values from 2 to 20 = −0.002, sd = 0.003). Finally, analogous cluster centroids were correlated between sets (**Figure S3, Figure S7**).

##### Feature-Free Clustering

To compare how the voxels cluster independently of their feature tunings, we followed a similar procedure to group voxels by their response profiles. Here, instead of using feature weights, we performed the clustering over voxels’ response magnitudes to each action video. All other procedures were the same as those used for feature-based clustering.

#### Preference Mapping Analyses

##### Comparing Gist and Body-Part-Action-Target Features

To compare performance based on the low-level gist model to the higher-level body part and action target models across visual cortex, we calculated a two-way preference map. First, both the gist model and the body-part-action-target model were used to predict responses in within-sets reliable voxels. Then, in each voxel, we found the model with the maximum *r*_CV_ and colored it so that the hue corresponds to the winning model and the saturation corresponds to the size of the difference between that value and the alternative model’s performance (e.g., a voxel where the gist model did markedly better than the body-part-action-target model is colored intensely yellow; **Figure 6b**).

##### Comparing Feature Involvement and Visibility

To compare models based on the functional involvement and visibility of body parts and action targets, we calculated a two-way preference map. As described above, each voxel was colored according to the model that predicted its responses the best (purple: feature involvement, blue: feature visibility; **Figure 7b**) and its saturation reflected the strength of the preference.

## ACKNOWLEDGEMENTS

Funding for this project was provided by NIH grant S10OD020039 to Harvard University Center for Brain Science, NSF grant DGE1144152 to L.T., and the Star Family Challenge Grant to T.K.

## AUTHOR CONTRIBUTIONS

L.T. and T.K. designed research; L.T. performed research; L.T. and T.K. contributed analytic tools; L.T. and T.K. analyzed data; L.T. and T.K. wrote the paper.

## DATA AVAILABILITY

Group fMRI and ratings data are available at the Open Science Framework repository for this project (https://osf.io/uvbg7/).

## CODE AVAILABILITY

Code for the main analyses is available at the Open Science Framework repository for this project (https://osf.io/uvbg7/).

## Supplemental Information

### Other supplementary materials for this manuscript include the following

Example frames for all stimuli, all feature ratings, pre-processed group fMRI data, and main analysis code for this paper are available at the Open Science Framework repository for this project (https://osf.io/uvbg7/). Single-Subject fMRI and ratings data available upon request.

## EXTENDED METHODS

### fMRI Data Acquisition

Imaging data were collected using a 32-channel phased-array head coil with a 3T Siemens Prisma fMRI Scanner at the Harvard Center for Brain Sciences. High-resolution T1-weighted anatomical scans were acquired using a 3D MPRAGE protocol (176 sagittal slices; FoV = 256 mm; 1×1×1 mm voxel resolution; gap thickness = 0 mm; TR = 2530 ms; TE = 1.69 ms; flip angle = 7 degrees). Blood oxygenation level-dependent (BOLD) contrast functional scans were obtained using a gradient echo-planar T2* sequence (84 oblique axial slices acquired at a 25° angle off of the anterior commissure-posterior commissure line; FoV = 204 mm; 1.5×1.5×1.5 mm voxel resolution; gap thickness = 0 mm, TR = 2000 ms; TE = 30 ms; flip angle = 80 degrees; multi-band acceleration factor = 3).

### fMRI Analysis and Pre-Processing

Functional data were pre-processed using Brain Voyager QX software version 2.8.4 (Brain Innovation, Maastricht, Netherlands). Functional preprocessing included slice scan-time correction, 3D motion correction, linear trend removal, temporal high-pass filtering (0.008 Hz cutoff), spatial smoothing (4 mm FWHM Kernel), and a transformation to Talairach coordinates. Whole-brain random-effect group GLMs were fit separately for each video set, as well as for both odd and even runs of each video set. In all cases, the design matrix included regressors for each condition block, specified as a square-wave regressor for each 5-second stimulus presentation time, convolved with a 2-gamma function that approximated the idealized hemodynamic response. Across these GLMs, the average variance inflation factor across conditions of the design matrix was 1.03 (where a value greater than 5 is considered problematic), and the average efficiency was 0.2 (Liu, Frank, Wong, & Buxton, 2001). Voxel time series were normalized within a run using a z-transform and corrected for temporal autocorrelations during GLM fitting. Beta weights extracted from these group-level random-effects GLMs were averaged across subjects for each voxel, and then taken as the primary measure of interest for all subsequent analyses. Each subject’s cortical surface was reconstructed from the high-resolution T1-weighted anatomical scan using Freesurfer software, and one subject was selected as the display brain for the group data.

### fMRI Experiment Design Details

Participants completed 8 functional runs of the experiment. Each video set was presented in four separate 6.2-minute runs. During each run, participants saw all 60 videos from one of the two sets. Each 2.5-second video was presented twice in a row, fading in and out of a uniform gray background over a 500-millisecond time window at the onset and offset of each presentation to prevent visually-jarring transients between video presentations. Thus, each video was presented in a 5-second block. In addition, four 15-second blocks of fixation were interspersed throughout the run, placed so that no fixation blocks occurred within five blocks of each other or the beginning or end of the run. In addition, fixation periods occurred for 4 seconds at the beginning and 10 seconds at the end of the run. Across runs, the order of the video blocks was randomized. Video stimuli were presented at 512 × 512 px on a 41.5 × 41.5 cm screen, subtending approximately 9 × 9 degrees of visual angle in the participant’s visual field. To ensure that participants remained alert throughout the experiment, they pressed a button whenever a red frame appeared around a video during one of the two video repetitions within a condition block. Such probes occurred 15 times per run and were counterbalanced so that each condition was probed once across all runs. All experimental protocols were presented using the Psychophysics Toolbox version 3 (Kleiner, Brainard, Pelli 2007) and MATLAB version R2016a.

### Orthogonalizing the Feature Spaces

To facilitate our interpretation of each feature’s contribution to the encoding models, the body parts and action target feature spaces were submitted to Principle Components Analysis (PCA), which extracts orthogonal components from the original feature space. Doing so enables weights to be fit over the body-part synergies rather than the body parts themselves – otherwise, with some kinds of regularized regression, highly correlated features (such as the thumb and index finger) might be assigned different and more variable weights, leading to the mistaken interpretation that a given region was tuned to the thumb but not the index finger. However, it is important to note that this step is not strictly necessary when using encoding modeling with ridge regularization. Therefore, it is equally statistically valid to skip this orthogonalization step and perform ridge-regularized encoding modeling over the original feature space.

PCA was performed separately on each feature space. The number of principle components (PCs) extracted was based on the number that cumulatively accounted for 95% of the variance in the feature ratings. This resulted in 7 body part PCs and 5 action target PCs (**Figure S1**). The encoding modeling analysis was then performed over these 12 PC features. Note that using these orthogonal predictors did not dramatically change the overall prediction performance compared to a model fit on the original 25 features (average change in cross-validated *r* = 0.02, sd = 0.11 for set 1; average change = 0.01, sd = 0.11 for set 2). We followed a similar approach to fit a model based on the body parts and targets that were *visible* in the videos. These ratings were averaged across raters for each video, then binarized by rounding the average rating to either 0 or 1. PCA was then performed separately on visibility ratings for body parts and action targets. Based on the number of PCs that cumulatively accounted for 95% of the variance in the feature ratings, 11 body part visibility PCs and 4 target visibility PCs were extracted. The encoding modeling analysis was then performed over these 15 PC features.

### Single-Subject Analyses

We also performed the encoding modeling and data-driven clustering analyses in each individual subject’s data. Because subjects vary in the extent of their reliable coverage, making comparisons across subjects challenging, encoding modeling analyses were done in the same voxels as in the group (cross-sets reliability > 0.30 in the group data). Similarly, voxels were clustered into five networks using the voxels that were reliable and well-predicted (*r*_CV_ > 0 with fdr-corrected q < 0.01) in the group data. To make it possible to compare the clustering results across subjects, in **Figure S4** the clusters are displayed on individual subjects’ brains using a common colormap: clusters with a similar tuning profile across subjects are displayed in similar colors. This colormap was made by entering all subject’s tuning profiles for all networks into a multi-dimensional scaling (MDS) analysis. The first three dimensions of the MDS solution were then extracted and used as R, G, and B values for each cluster’s display color. Because some subjects did not have reliable neural responses in all of these regions, their results in **Figure S2** and **Figure S4** are ordered by the number of reliable voxels found in each subject’s brain when reliable voxels were defined individually for each subject (cross-sets reliability > 0.30 in each subject’s data).

### Hierarchical Clustering

To validate our finding (based on *k*-means clustering) that the division between regions tuned to social and non-social features of actions emerges first, we also conducted a hierarchical clustering analysis. Voxels were grouped into a hierarchical tree based on the similarity of their feature weight profiles (the weights assigned by the voxel-wise encoding model to the 12 body-part and action-target features). This analysis was conducted separately over the data from our two video sets, using the group data. The analysis was restricted to the same set of voxels used in the k-means clustering analysis: voxels that were reliable across video sets and predicted well (*r*_CV_ > 0 with q < 0.01 after FDR-correction) by the model when it was fit to the data from either video set. Next, we used MATLAB’s linkage function with the correlation distance metric to cluster voxels and then arrange them into a hierarchy based on the average distance between clusters. To determine the predominant division within this hierarchy, we examined the results at the point where the voxels branched into two clusters (**Figure S6**). To determine the robustness of this clustering, we computed d-prime between the voxel groupings for video sets 1 and 2. To determine how similarly voxels were grouped by hierarchical clustering and *k*-means clustering, we computed d-prime between the voxel groupings for the two methods, within each video set. Additional analyses revealed that subsequent divisions found by the hierarchical clustering analysis formed clusters that were very small (e.g., 30 voxels or fewer), and thus this supplemental analysis was not pursued further.

### Relating Motion Span and Interaction Envelope

The span of motion present in each action video was measured in two ways. First, human raters (*N* = 182) completed an online experiment on Amazon Mechanical Turk, in which they watched each video and then answered the question, “How much movement does this action or activity involve for the average person?” using a 1 to 5 scale. Second, optical flow was calculated for every frame in the videos using the Horn-Schunck algorithm, implemented in MATLAB (Horn & Schunck, 1981). The proportion of pixels that contained movement (average optical flow magnitude > 0.01) was then calculated for each video. To relate these measures of movement to neural tuning patterns, we first obtained the overall neural response to each video for each of the five networks, averaging across all voxels in each network. Next, we calculated each network’s motion sensitivity using a weighted average measure (i.e. for each network, the beta estimate for each video was multiplied by that video’s motion span; these products were then summed together and divided by the total number of videos). Motion sensitivity is therefore an average response across all the videos, weighted by the amount of movement in each video. Each network’s motion sensitivity was calculated separately for the two types of movement measurements (human ratings and optical flow; **Figure S8**). This analysis was done separately for the two video sets.

### Sociality and Transitivity Analysis

To compare our data to results reported in Wurm & Caramazza (2017), we searched for regions that were preferentially tuned to sociality (directed at a person) or transitivity (directed at an object). To do so, we fit a separate encoding model based on the raw, un-PC’d action target feature matrix. Then, in each voxel we compared the magnitude of the weight assigned to object targets with the weight assigned to person targets. Voxels were colored orange if the object weight was larger and pink if the person weight was larger. Saturation reflects the size of the difference between the weights (**Figure S9**).

---

**Figure S1.**
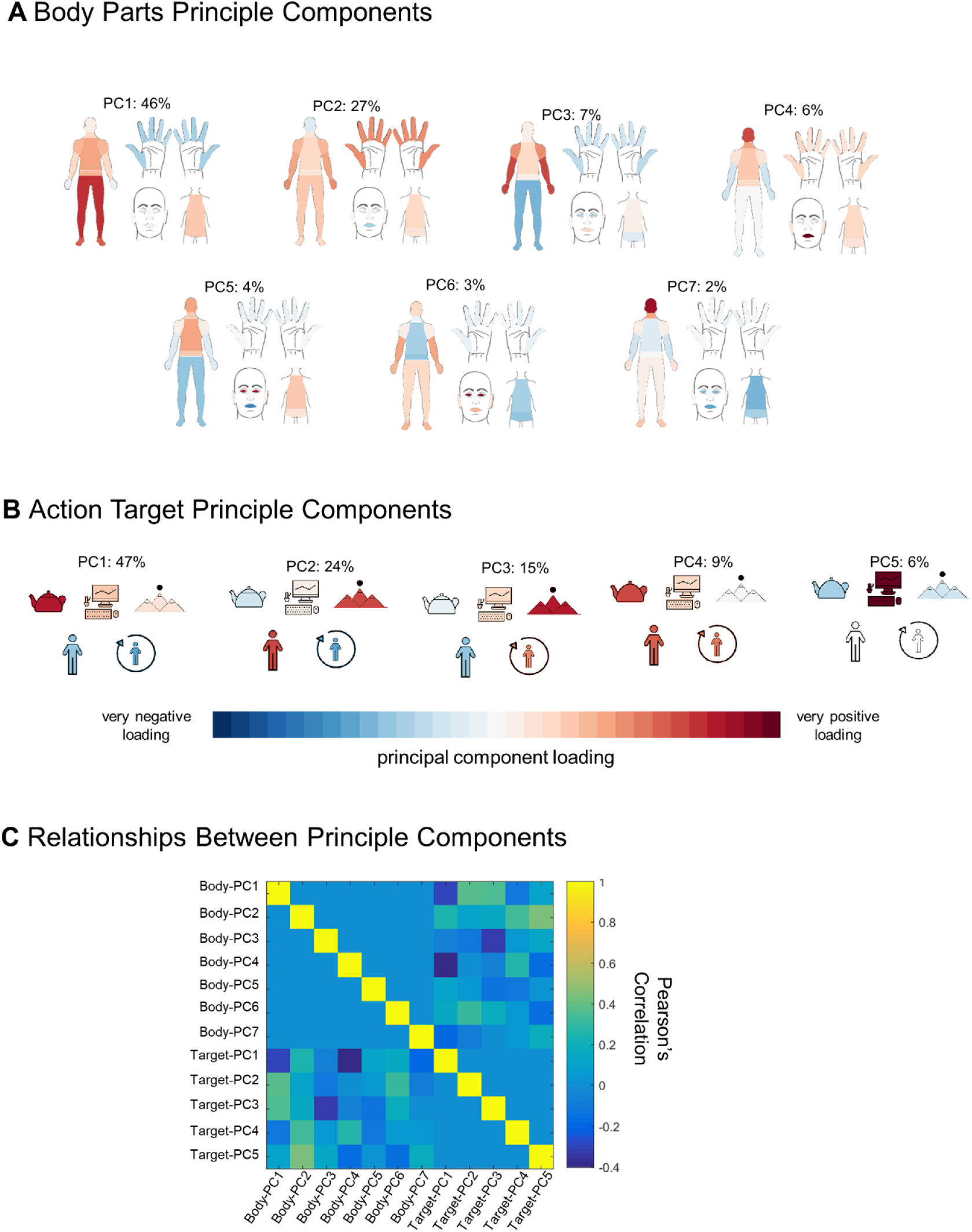
Principle Components Analysis of Body Part and Action Target Feature Spaces. Visualization of the (A) body parts and (B) action target features after being reduced via Principle Components Analysis. Percentages indicate percent variance in the feature ratings explained by each component. (C) Relationships between the Principle Components features, measured using Pearson’s correlation.

**Figure S2.**
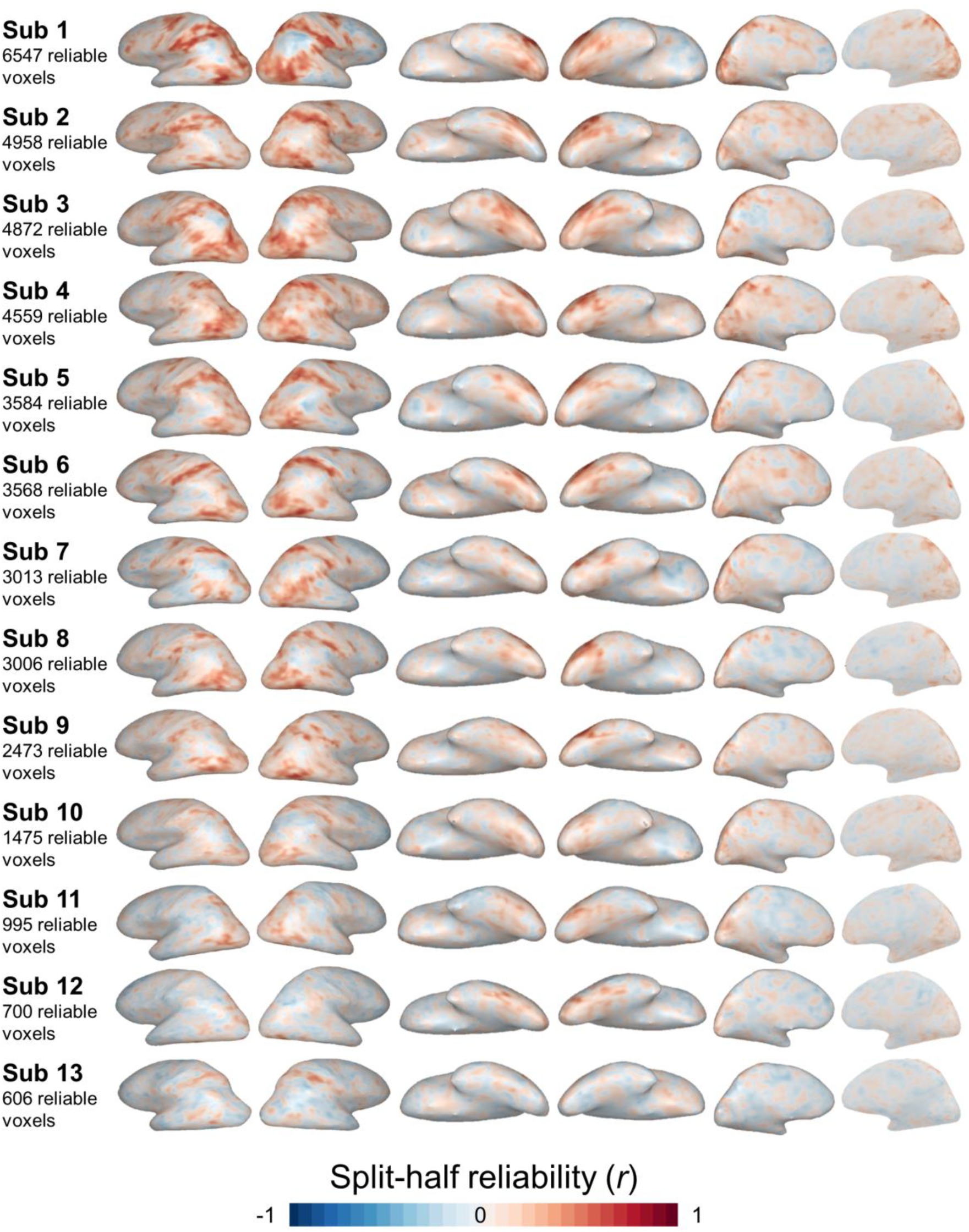
Voxel-wise Reliability in Individual Subjects. Split-half reliability maps are shown for each subject. Subjects are ordered according to the number of voxels that survived the reliability-based inclusion threshold (split-half *r* > 0.30, which was an appropriate threshold for all subjects, as well as in the group data).

**Figure S3.**
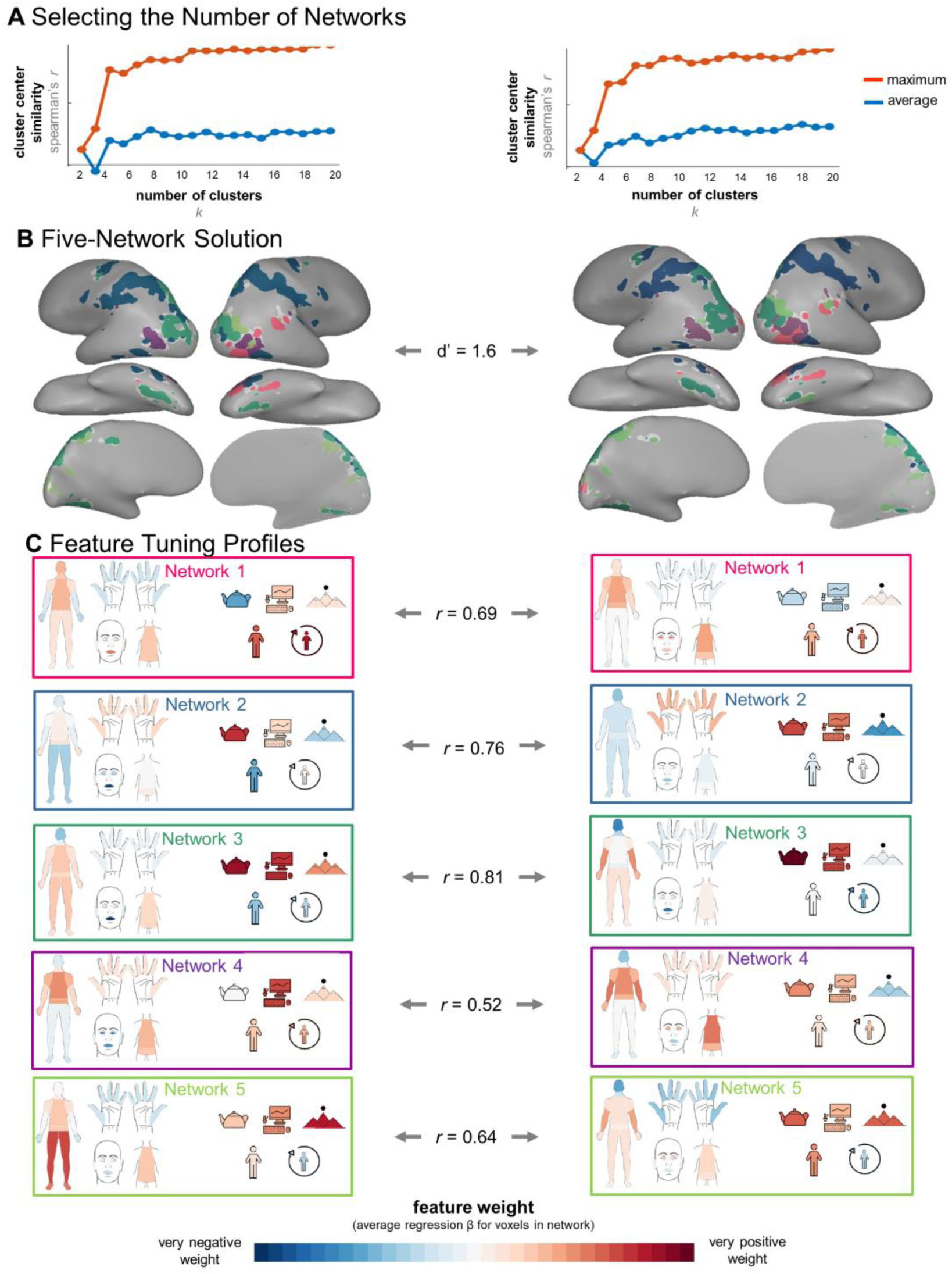
Comparing 5-Network Solution Across Stimulus Sets. Results are shown for video set 1 (left) and set 2 (right). (A) *K*-means clustering was performed at every *k* from 2 to 20 (x-axis), and the resulting cluster centroid similarities (measured as the average (blue) and maximum (orange) correlation between the centers of every cluster) are plotted. (B) The 5-network structure is displayed for each video set. The match between the voxel assignments was computed using d-prime. (C) The feature tuning profile is shown for each cluster. Tuning profiles were compared across video sets using Pearson’s correlation.

**Figure S4.**
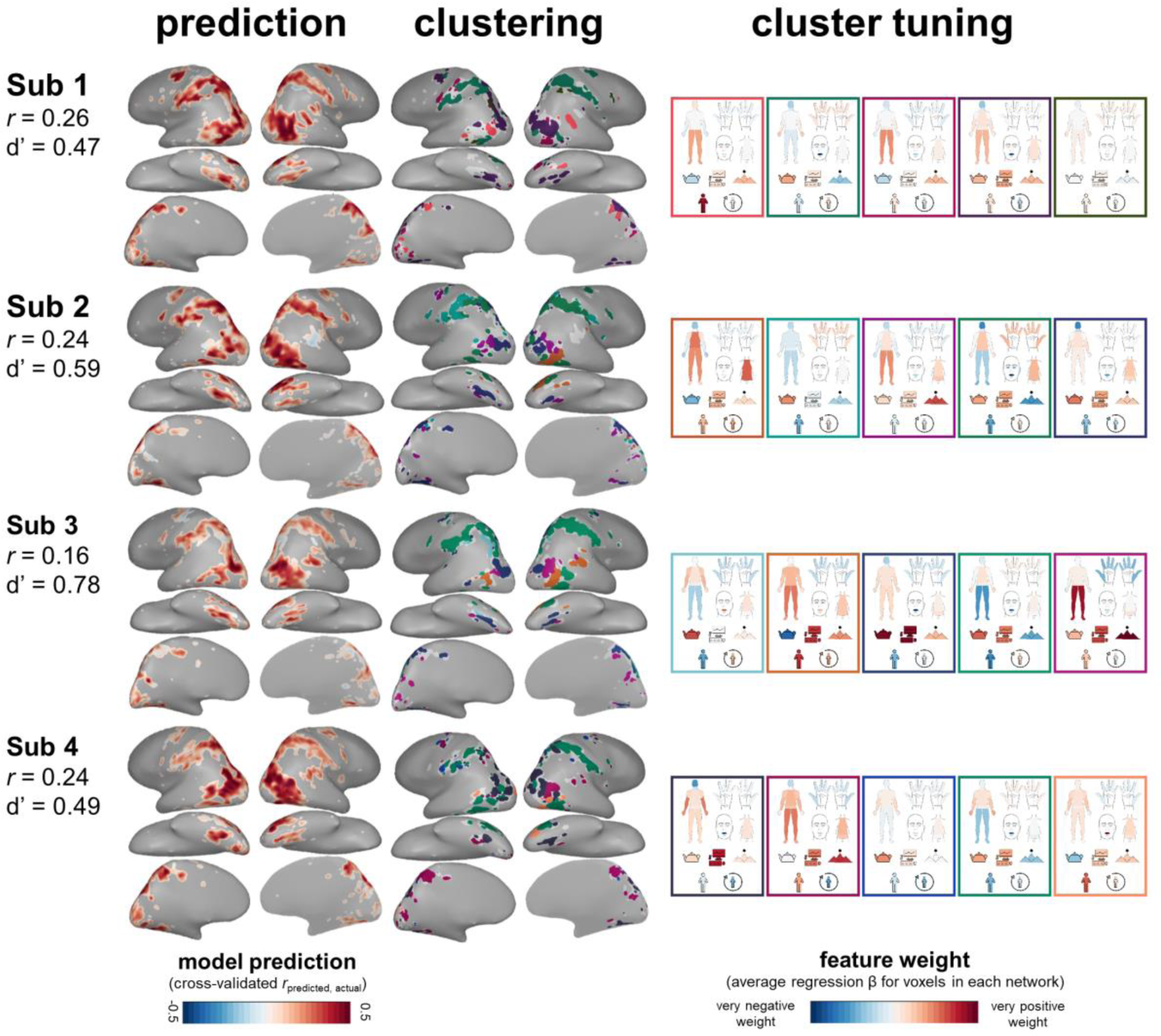
Model Performance and Large-Scale Structure in Individual Subjects. Prediction performance for the action-target-body-part model and large-scale clustering of the model weights into five clusters are shown for each subject. For clustering results displayed on the brain, voxels are colored according to a common colormap: all voxels in
a cluster are shown in the same color, and clusters with similar tuning profiles are shown in similar colors. In addition, each cluster’s tuning profile is displayed to show which features the clusters are sensitive to. For each subject, we also list the median crossvalidated prediction performance (*r*) and d-prime comparing how voxels were grouped in the single-subject and group data. All results are based on the data from video set 1. Subjects are ordered according to the number of voxels that survived the reliability threshold in their data (see **Figure S2**). All single-subject analyses were conducted in the voxels used in the group analysis, though reliable coverage within these voxels differed across subjects. Figure continues for Subjects 5-13 on the next two pages.

**Figure S5:**
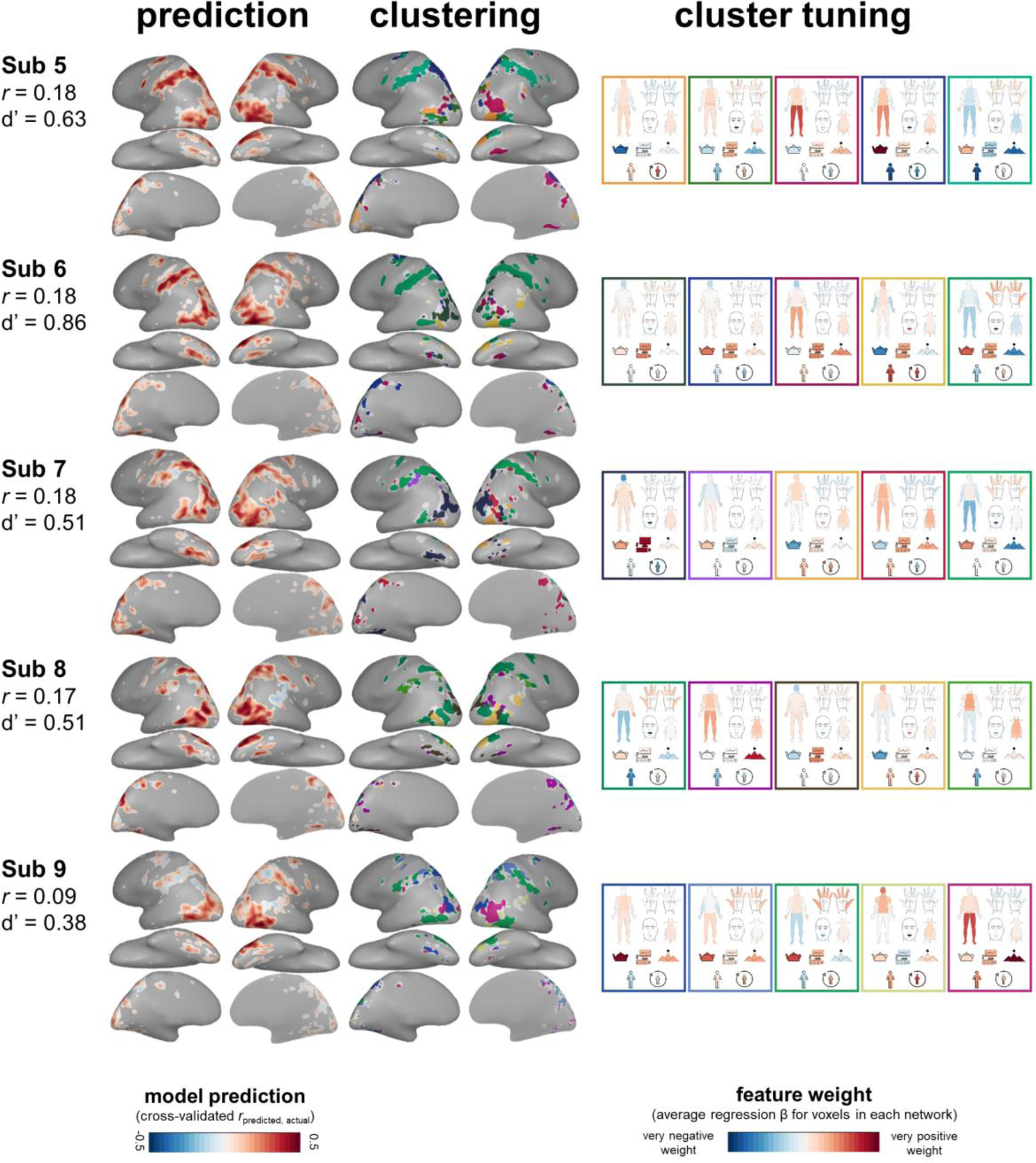
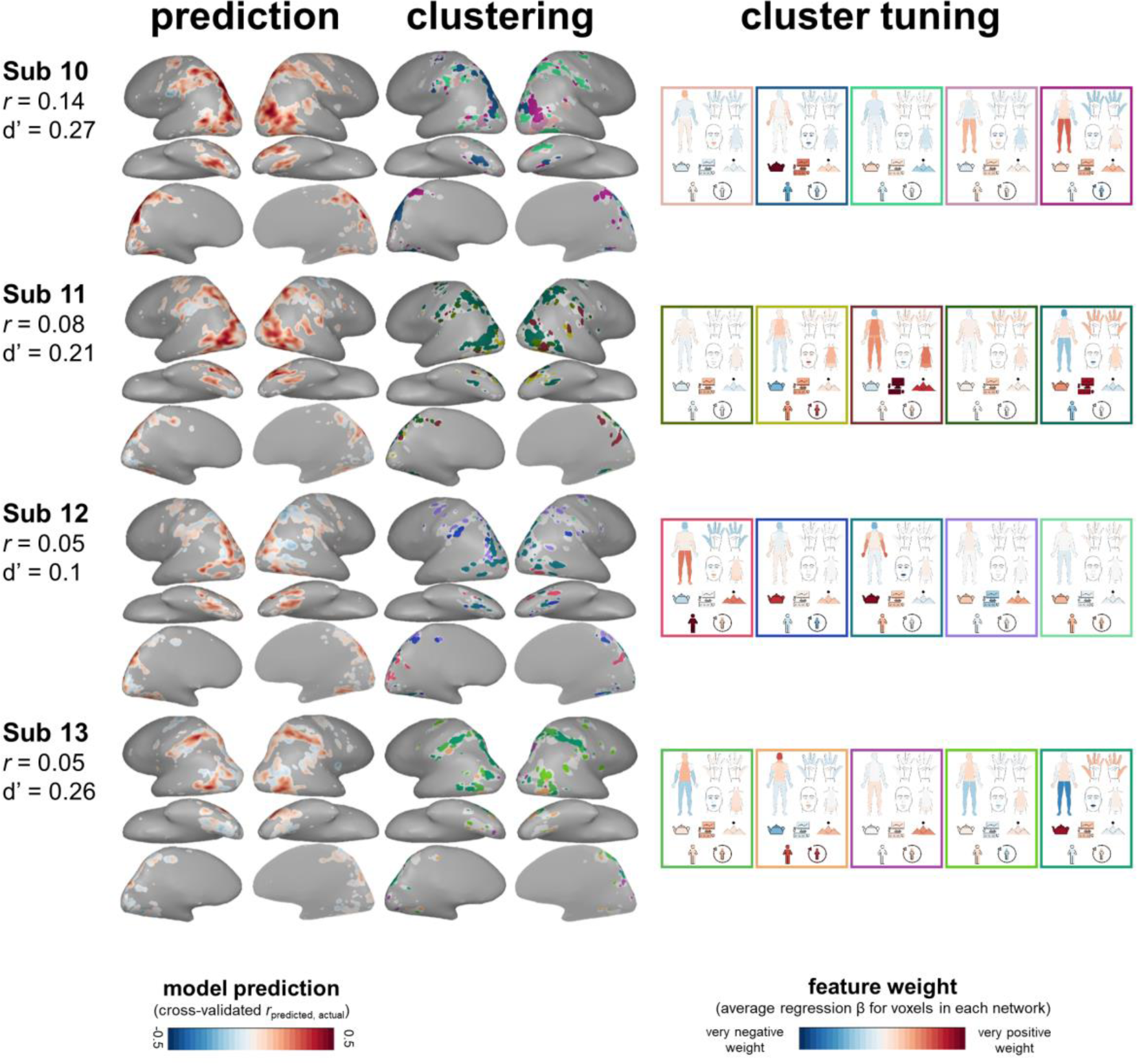
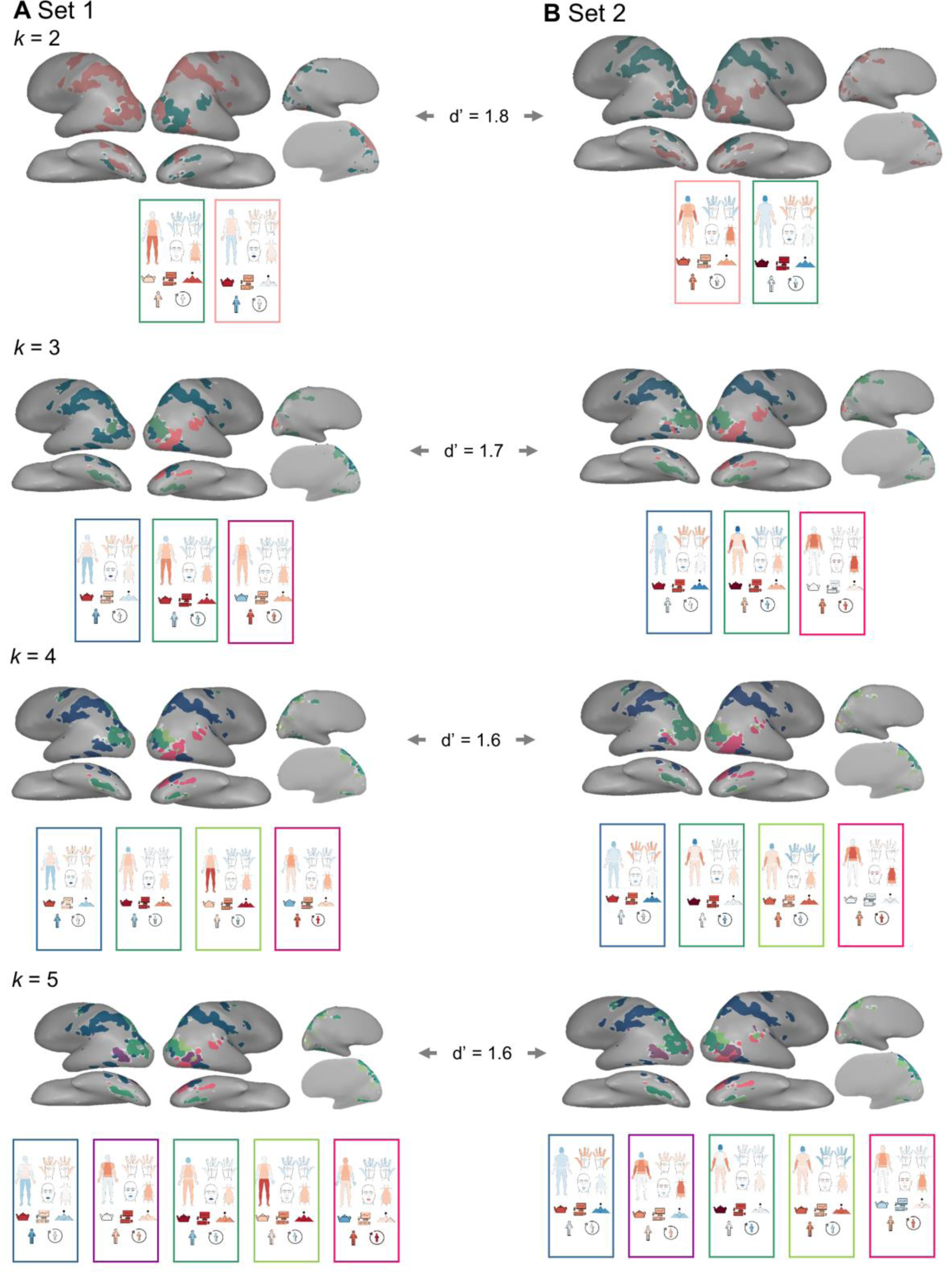
The Emergence of Five Large-Scale Networks. Clustering solutions are shown at *k* = 2, 3, 4, and 5 clusters for (A) video set 1 and (B) video set 2. D-prime values indicate the correspondence between how voxels were clustered across the video sets.

**Figure S6:**
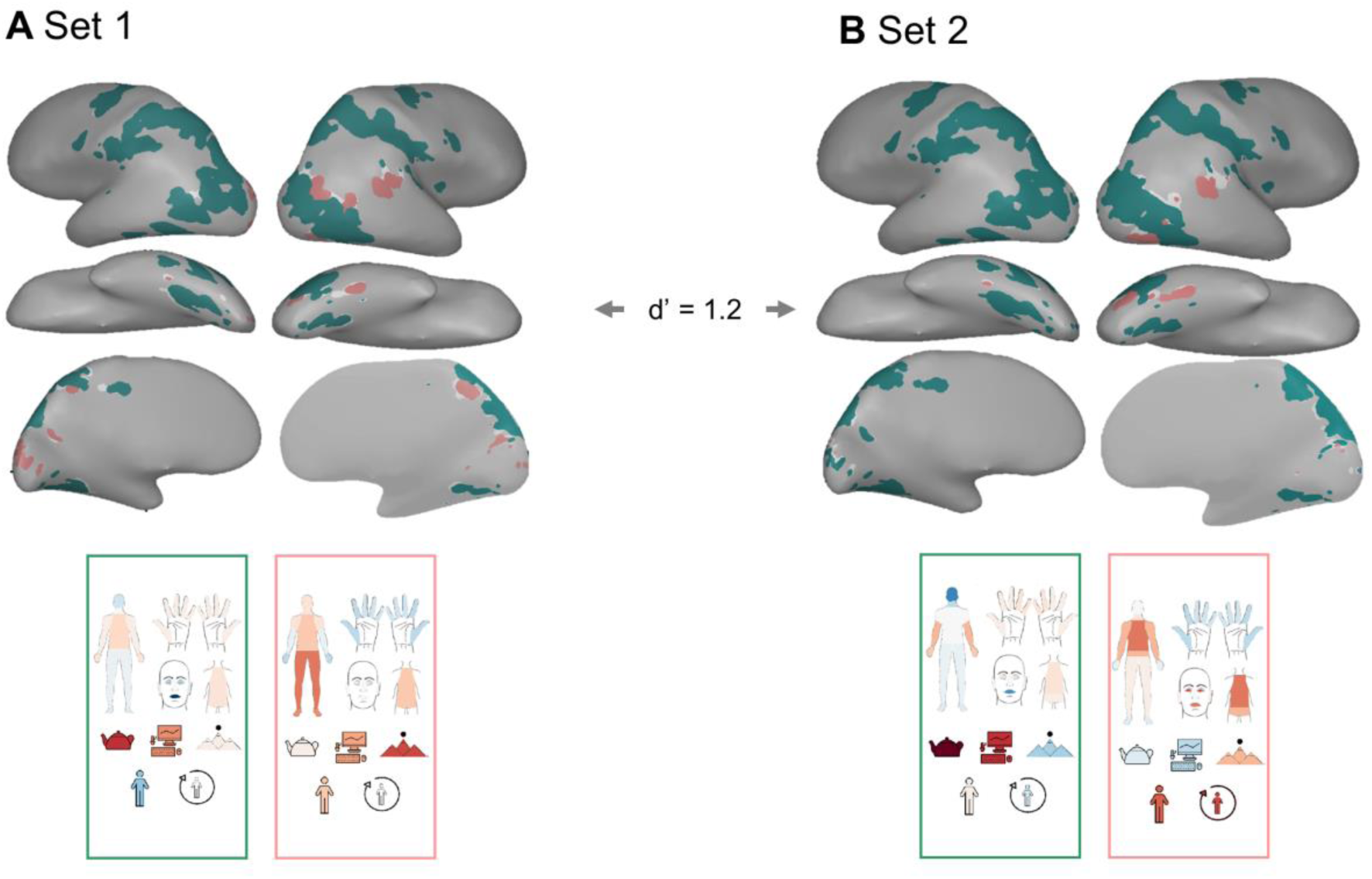
Hierarchical Clustering of Voxel-wise Feature Tuning. Results of hierarchically clustering voxels into 2 clusters based on their feature tuning are shown for (A) video set 1 and (B) video set 2. The feature tuning profile (the average tuning across all voxels in the cluster) is shown for each cluster. D-prime value indicates the correspondence between how voxels were clustered across the video sets. The voxels were grouped similarly using this method compared to k-means clustering at *k* = 2 (**Figure S5**; d-prime across clustering methods = 1.6 (set 1) & 0.6 (set 2)).

**Figure S7.**
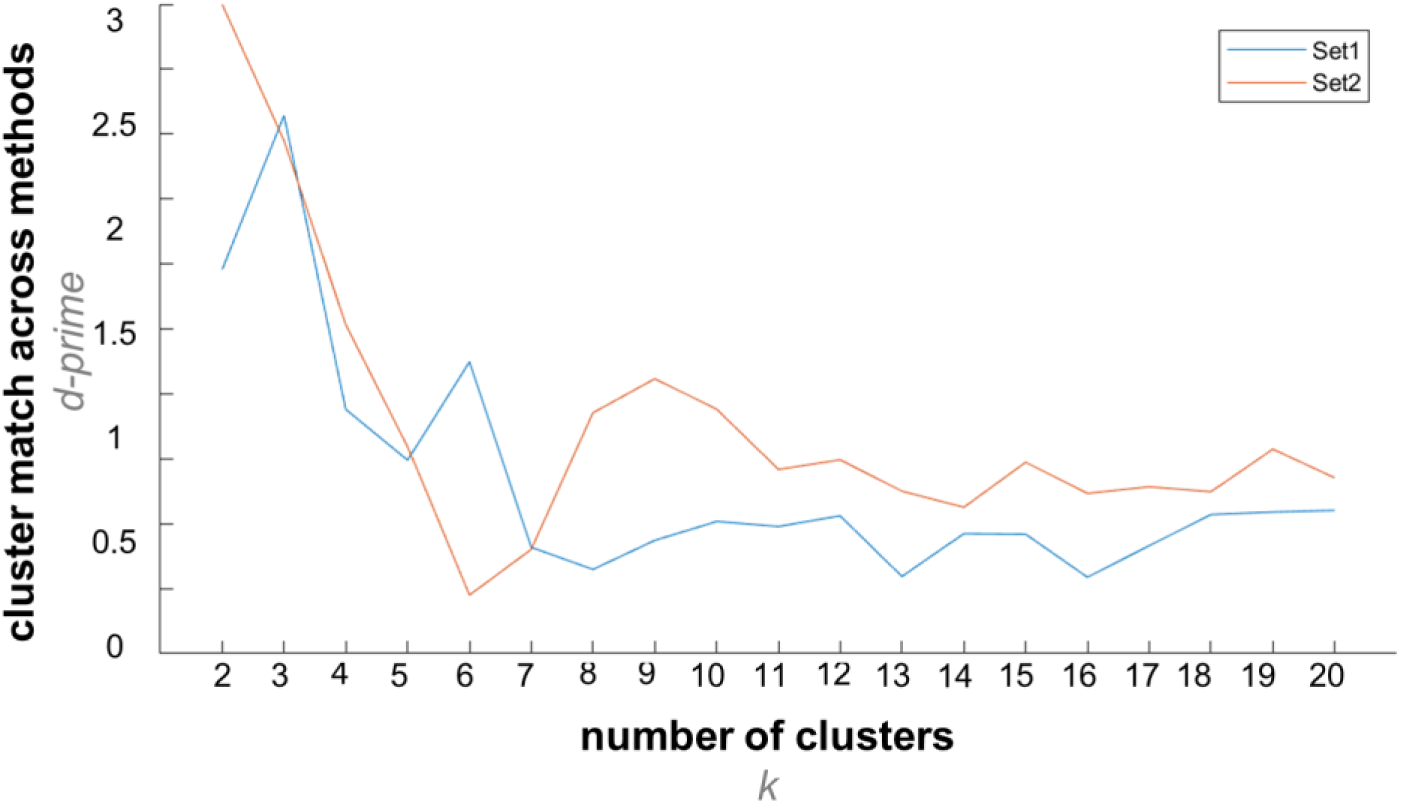
Comparing Feature-Based and Feature-Free Clustering Analyses. Feature-free and feature-based clustering solutions were compared at a range of possible numbers of clusters (*k*-values) using d-prime. This was done separately for each video set.

**Figure S8.**
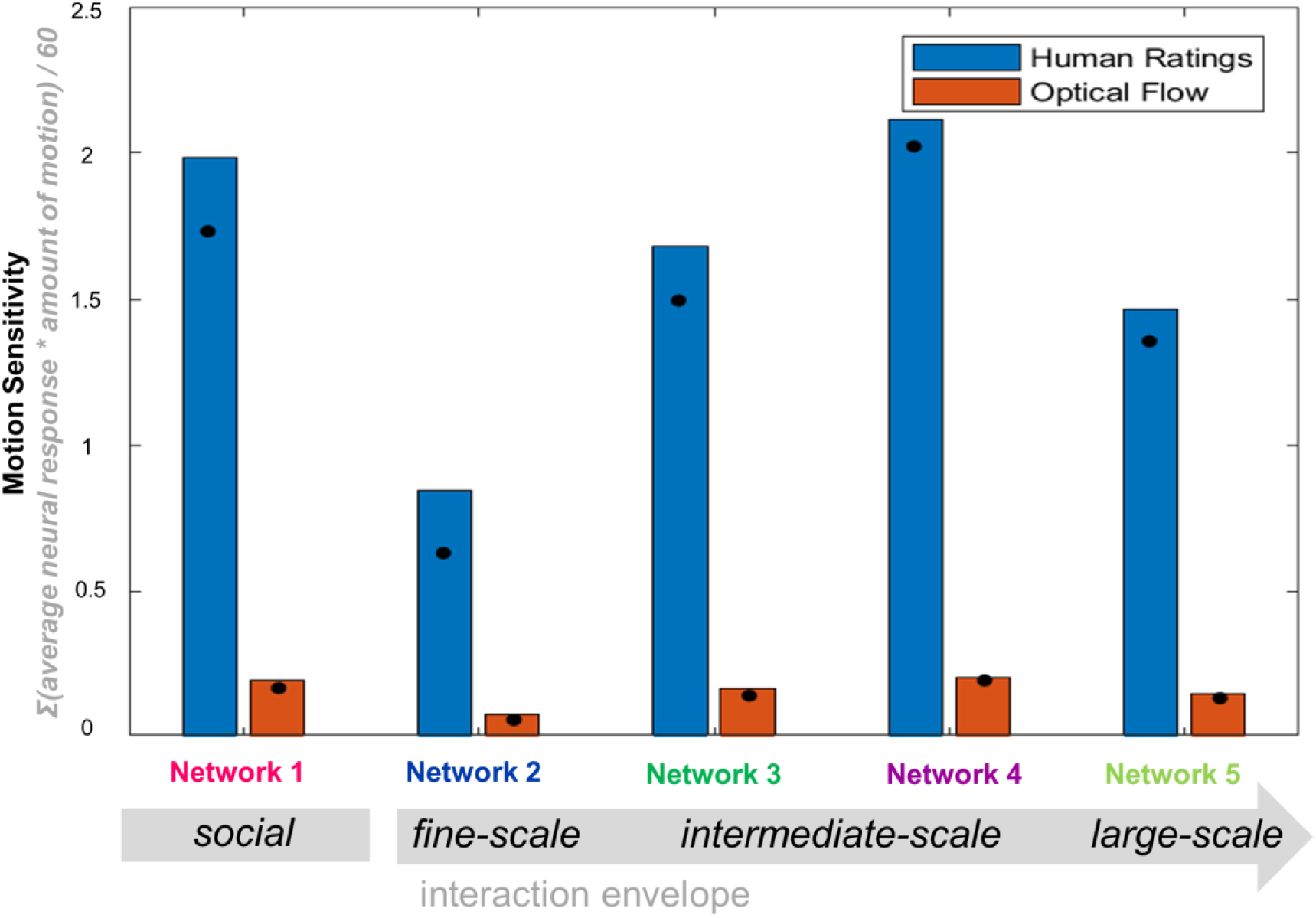
Relating Interaction Envelope and Motion Span. Sensitivity to the span of motion in the action videos is plotted for the five action sub-networks. Motion Sensitivity was calculated as each network’s average response over the action videos, weighted by the motion span apparent in each video. Blue bars depict motion sensitivity in video set 1 based on human ratings of how much the average actor moves to complete each action. Orange bars depict motion sensitivity in video set 1 based on the proportion of pixels in the video’s frame containing movement during the course of the video, measured by optical flow. Black dots indicate the same values for video set 2.

**Figure S9:**
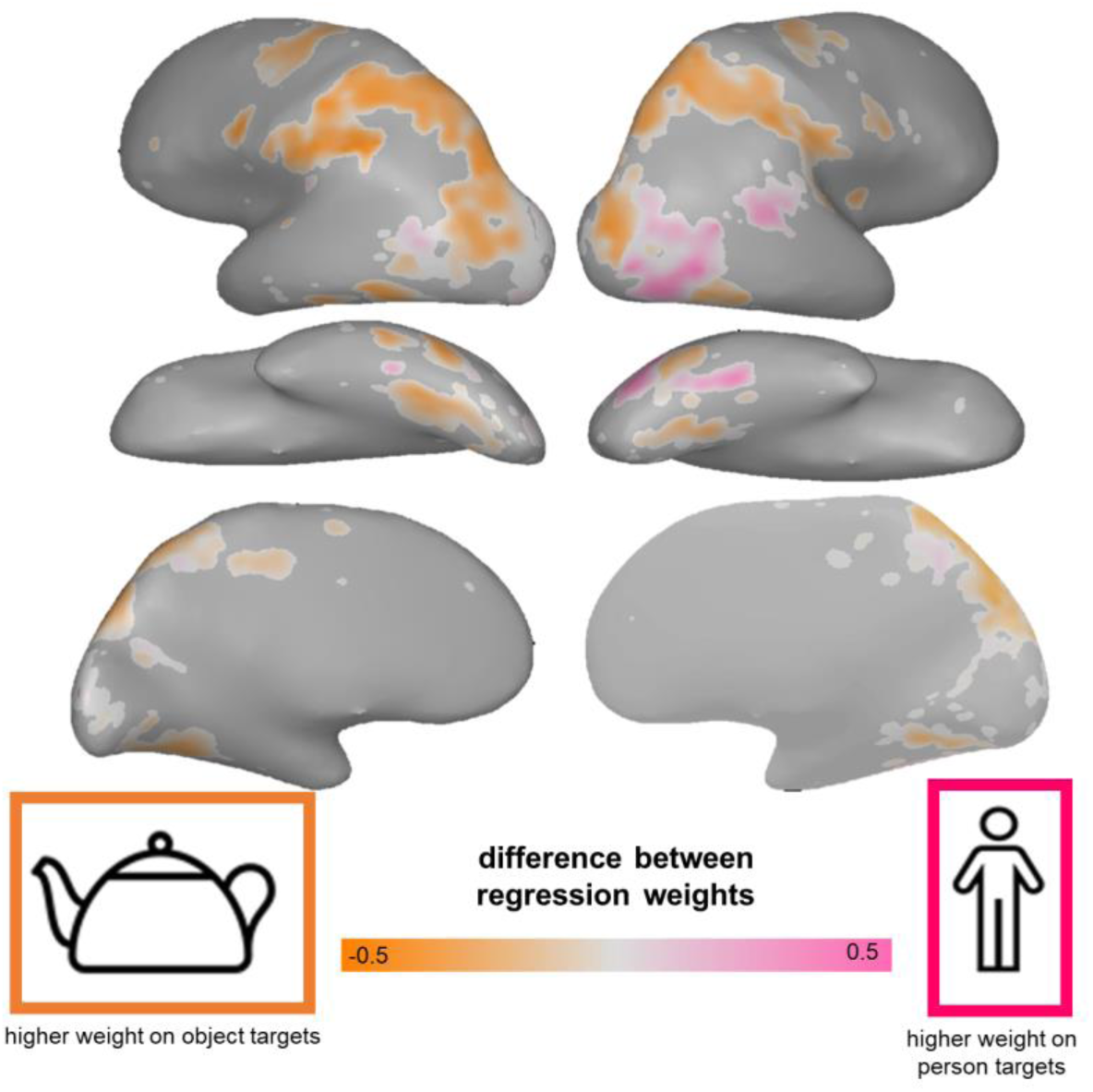
Sociality and Transitivity (alternate feature spaces). Two-way preference map showing regions more related to person targets (“social” analog) or object targets (“transitive” analog). Voxels are colored according to the feature with the highest weight of the two possible targets: pink for person > object, orange for object > person. Color saturation reflects the strength of the voxel’s preference (person weight – object weight).

